# Breaking β-sheets in FUS prion-like domain preserves phase separation and function but prevents aggregation and toxicity

**DOI:** 10.64898/2026.02.17.706410

**Authors:** Noah Wake, Juan Alcalde, Daniel Jutzi, Anjali Bajaj, Sukhleen Kour, Mayur Barai, Shuo-Lin Weng, Samara Cummings, Tongyin Zheng, Eric N. Anderson, Szu-Huan Wang, Ryan Puterbaugh, Daryl A. Bosco, Benjamin S. Schuster, Jeetain Mittal, Udai Bhan Pandey, Marc-David Ruepp, Nicolas L Fawzi

## Abstract

The RNA-binding protein Fused in Sarcoma (FUS) undergoes phase separation associated with RNA processing. However, the prion-like low complexity (LC) domain of FUS forms solid-like aggregates in neurodegenerative diseases. Whether the formation of β-sheet structure associated with pathology is also physiologically/functionally relevant is debated. Similarly, if mislocalization alone or concomitant aggregation is responsible for FUS gain-of-function toxicity remains to be probed. Here, we introduce β-sheet breaking proline residues into FUS LC with the goal of preventing cross-β-driven aggregation without disrupting essential functions and phase separation. β-sheet-deficient FUS variants maintain native-like global motions, disorder, and phase separation, but no longer show a liquid-to-solid transition (LST). Biochemical partitioning, cellular localization, and auto- and cross-regulatory functions of FUS all remain essentially unchanged. Conversely, FUS-induced neurodegeneration in several *Drosophila* models is drastically reduced. These findings suggest a strategy for mitigating disease-related toxicity through backbone structure modulation to prevent prion-like domain protein aggregation.

**GRAPHICAL ABSTRACT:** 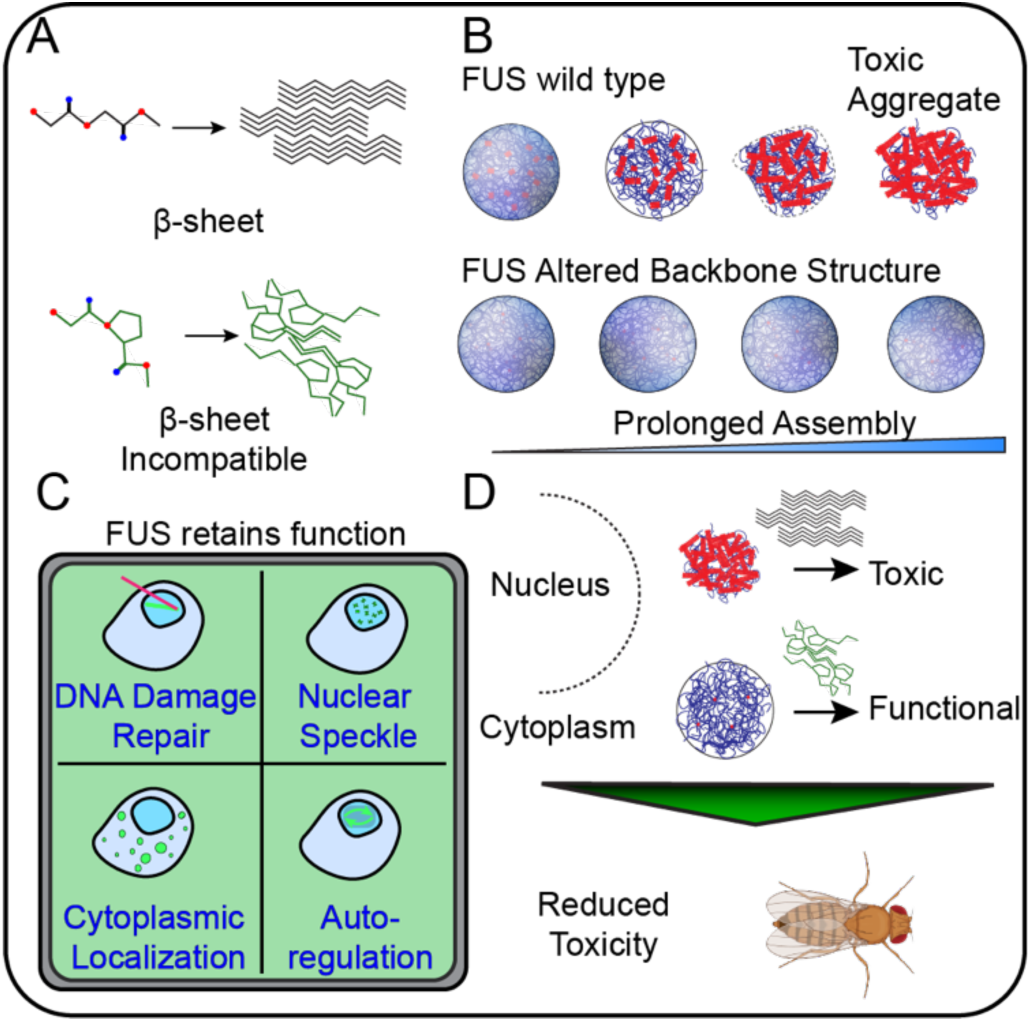

**SUMMARY:** The RNA-binding protein Fused in Sarcoma (FUS) undergoes phase separation as part of its physiological function but can aberrantly aggregate into solid-like assemblies in amyotrophic lateral sclerosis and frontotemporal dementia. To dissect the role of β-sheets in both function and pathological transition, we engineered β-sheet-preventing FUS variants via targeted proline residue insertions in the prion-like disordered region. These variants retained native structure, motions, and phase behavior yet showed dramatically reduced aggregation, both as an isolated prion-like domain and in full-length FUS. Crucially, these variants maintained a panel of FUS cellular functions that depend on FUS condensation but prevented FUS toxicity in fly models of neurodegeneration. Our findings implicate β-sheets as key drivers of FUS condensate maturation and neuronal toxicity, highlighting β-sheet modulation as a therapeutic strategy against FUS-related neurodegeneration.

**HIGHLIGHTS:**

- Targeted proline additions disrupt β-sheet formation in FUS without altering native conformations, dynamics, or phase separation behavior
- β-sheet-deficient FUS variants prevent aggregation and liquid-to-solid transitions while retaining key biological functions
- In vivo models reveal attenuated toxicity of β-sheet-deficient FUS in *Drosophila*
- β-sheets are identified as central drivers of condensate maturation and neuronal death, offering a therapeutic entry point for modulating prion-like domain pathology

## Introduction

Phase separation into liquid-like compartments including ribonucleoprotein granules is an important feature of several RNA-binding proteins^1,2^. Fused in Sarcoma (FUS) is a key constituent in functional biomolecular condensates as well as in pathological inclusions in amyotrophic lateral sclerosis (ALS) and frontotemporal lobar degeneration (FTLD)^3–5^, yet how it transitions from dynamic, liquid-like assemblies to solid-like aggregates, and how this transition contributes to FUS gain-of-function toxicity remains poorly understood. Despite the proposed connection between solid fibrillar forms of FUS and disease, the liquid-to-solid transition (LST) of other condensate-forming proteins can have a functional role^6^. Thus, to understand the factors that modulate the diverse material properties ranging from liquid-like to solid-like and their connection to function and toxicity, it is essential to probe the structural features that facilitate the LST and to correlate these structures with FUS toxicity.

In healthy cells, RNA metabolism^7^ and the integrated stress response^8^ rely on the formation of biomolecular condensates^1^. Several studies have demonstrated that FUS phase separation and function depend on its low-complexity (LC) domain^9,10^, which shares the sequence enrichment of polar amino acids with yeast prion proteins^11,12^. Similar to how the primary sequence of globular proteins directly determines their three-dimensional structure and function, rules governing the relationship between primary sequence composition (especially the important role of aromatic and arginine residues) and liquid-like phase separation have been demonstrated^13,14^. Further studies expanding the molecular grammars of phase separation have demonstrated the role of hydrophobic and polar residues^14–18^. Combined with studies showing that these domains remain disordered in condensates^19^, there is strong support for a model where phase separation of low complexity prion-like domains involves multivalent interactions between structurally disordered regions^15,20^. Going further, recent studies using FUS as a model have shown how local sequence context can also modulate residue-type contributions to aggregation^14^. However, these studies have not directly addressed a long-standing debate over whether the interactions driving cellular functions associated with phase separation occur between segments that remain disordered as envisioned in the models above^15,20,21^ or if these sequence regions function like short linear motifs (SLiMs)^22,23^ that can fold into Long Aromatic-Rich Kinked Segments (LARKS)^24,25^ or form other transient/labile β-sheet interactions^26,27^ to stabilize phase separation.

Beyond the structural details, understanding of how FUS self-assembly and interactions underlie its many nuclear and cytoplasmic functions that can become dysregulated in disease has continued to grow^28,29^. In the nucleus, FUS contributes to recruitment of RNA polymerase II (Pol II) to sites of transcription^30^ and alters gene expression by selectively partitioning genomic regions—“gene modules”—that are primed for transcriptional activation and co-expression^31^. In response to cellular stress, FUS can be exported to the cytoplasm, where it localizes in stress granules that sequester mRNAs and stalled pre-initiation complexes^32^. Upon resolution of the stress, FUS binds the nuclear import receptor protein transportin-1/karophyerin-β2 via a PY nuclear localization signal (NLS) to be shuttled back into the nucleus^33^. Indeed, many ALS-causing dominant mutations of FUS alter its nuclear localization by introducing nonsense (e.g. R495X), frameshifts (e.g. G515Vfs*14), or point mutations (e.g. P525L) that directly affect NLS recognition^33,34^, supporting a cytoplasmic gain-of-toxic function mechanism. While stress granules are important components of the integrated stress response and serve as a protective mechanism, it has been proposed that prolonged cytoplasmic localization may nucleate the formation of amyloidogenic inclusions that are associated with the toxicity observed in ALS and FTLD^35^. In FUS-ALS, wild-type FUS can also be mislocalized, suggesting a concomitant loss-of-nuclear-function along with gain-of-toxic-function is possible^36^, and together contribute to motor neuron loss. While the loss-of-function toxicity has been ascribed to the depletion of active populations of nuclear FUS, it is not well understood which disease-linked features trigger toxicity. While phase separation is recognized as essential for proper FUS function^37^, it remains unclear whether the liquid-to-solid transition plays a functional role in cellular processes. Some studies suggest that formation of reversible stress granules with “solid-like” material properties is crucial to the integrated stress response^38^, where heterogeneity in stabilizing interactions modulates condensate material properties. In hnRNPA2, reversible amyloid cores (RACs) have been proposed to regulate droplet fusion rates via stacking of charged residues in the fibril core^39^. Conversely, disease-linked mutations in hnRNPA2 that enhance β-sheet character disrupt this delicate balance by promoting the formation of β-rich, irreversible protein aggregates^40–42^. Simulation studies examining the effects of RNA on LST of FUS or hnRNPA2 suggest that the progressive accumulation of persistent β-sheet interactions – with contact lifetimes that must be orders of magnitude longer than transient interactions stabilizing phase separation – alters the density of aging condensates, ultimately leading to irreversible β-sheet aggregates^43^. Others suggest that β-sheet motifs can be altered by bulky residues or enhanced in disease mutations^40^ and that this may modulate the material properties of the condensate, suggesting a relationship between protein structure and condensate maturation. On the other hand, some studies claim that the short peptide regions that drive aggregation also facilitate liquid-like phase separation by forming labile interactions mediated by β-sheets^44,45^, though direct structural evidence supporting these models is lacking^15^. However, it remains untested whether these β-sheet structures play a functional role in FUS activity in cells, where it is difficult to directly visualize their structure, or solely promote the conversion to solid fibrils. Much of the gap arises because studies examining the role of domains like FUS LC that drive both phase separation and aggregation have done so by domain deletion or by mutations that disrupt both processes, such as changing several tyrosine residues to serine^37,44,46^. Thus, delineating and separating condensation, aggregation, and the LST – and probing their relationship to FUS function and toxicity – is critically important.

Recent experiments show that increasing glycine content delays FUS aggregation, suggesting that the greater conformational entropy penalty required for ordering the peptide backbone of glycine in a β-sheet discourages aggregation^14,47,48^. Although often thought of as “stiff” in contrast to “flexible” glycine, proline residues cannot form backbone hydrogen bonds needed in β-sheet structure and hence have also been shown to inhibit protein aggregation in several contexts^49–52,53,54,55^ including, in the case of TIA1, removal of native prolines speeds LST. In a therapeutic context, the substitution of residues with proline found at homologous positions in amylin sequences of other animals disrupts aggregation and was used to improve amylin-mimicking peptides^56,57^. Similarly, studies of synthetic peptides by Rauscher et al^58^, demonstrate that amyloid fibril formation is impeded above a threshold of sequence proline composition. Together, these findings suggest that both proline and glycine can play roles in preventing aggregation by discouraging β-sheet structure, further supporting a hypothesis in which alterations to protein backbone structure affect aggregation. However, the mechanistic basis of the effect of glycine and proline on aggregation in these studies and whether the β-sheet interactions stabilizing aggregates are similarly important for phase separation, function, and toxicity, is not well understood.

Despite extensive efforts to elucidate the molecular mechanisms underlying FUS’s phase separation and liquid-to-solid transition, the stages of the transition from liquid to solid in which FUS becomes toxic remain unclear. In this study, we probe how selective modifications to the backbone structure of the prion-like domain of FUS alter phase separation and/or the LST, elucidating if these are linked. Additionally, we probe if FUS β-sheet aggregates are critical for FUS functions in cells and if they contribute to FUS toxicity in *Drosophila* neurodegeneration models. Our studies aim to test if in fact β-sheet structure is required for FUS phase separation and/or toxicity. Importantly, these studies aim to test if prion-like domain phase separation/function and aggregation can be decoupled, which could then enable several therapeutic avenues given the prevalence of neurodegenerative diseases associated with aggregation of functionally essential proteins.

## Results

### FUS phase separation and aggregation occur through independent mechanisms

To investigate the relationship between phase separation and aggregation, we hypothesized that precisely modifying the sequence of the aggregation-prone, prion-like, low-complexity (LC) domain of FUS could selectively prevent aggregation while preserving phase separation. We targeted an amyloid core region that we recently showed was formed in test tubes^14^, as well as another reported fibril core^59^. Specifically, we aimed to engineer a FUS variant with reduced potential to form β-sheets that would allow us to disentangle the molecular determinants of interactions stabilizing phase separation, and potentially function, from aggregation. We targeted local sequence contexts associated with β-sheet structures such as steric zippers using ZipperDB^60^ as well as regions prone to β-arcs and superpleated β-sheet structures^61,62^ (**Fig. S1A**). Guided by these predictions, we inserted proline residues into regions with high β-sheet formation propensity, preserving the local sequence grammar by not removing any residues and retaining the SQ and TQ sites that are known targets of phosphorylation by DNA-dependent protein kinase^9^ (**Fig. 1A, Fig. S1A**). We created several different variants with increasing number of proline additions for comparison. Because of our recent finding that specific serines within a FUS fibril core significantly altered FUS LC aggregation^14^, we targeted specific β-sheet segments within the this core region, which we name FUS 4P as it adds four proline residues. Based on other stable amyloid core structures shown for FUS LC fibrils^59^ and the potential for fibril core rearrangement due to the mutations, we generalized this approach to extend beyond reported amyloid core structures formed *in vitro* and targeted sequence regions predicted to form additional β-sheets throughout the entirety of the FUS LC sequence (FUS 12P). Finally, we combined these two approaches (16P). The β-serpentine algorithm showed a reduction in predicted β-sheet formation correlated with increasing proline content for each of these variants with ∼95% reduction in total β-sheets for all three variants (**Fig. 1B**). To confirm that our engineered sequences retained their prion-like sequence features, we used the Toombs (PLAAC) algorithm to evaluate the sequence similarity to known yeast prions^51^. Although naturally occurring yeast prion domains are not significantly enriched in proline^51^, the prion-like sequence character of these variant remained nearly identical to that of wild-type FUS (**Fig. 1C**). We then compared the phase diagrams of these variants to understand what, if any, effect β-sheet regions had on phase separation. First, we observed that each variant retained the ability to form condensates that had morphologies consistent with condensed, liquid-like phases (**Fig. 1D**). Next to test the effect of the added prolines on the driving force for phase separation, we measured the saturation concentration of each variant under increasing concentrations of sodium chloride. We found that all three variants phase separated similarly in response to changes in the ionic strength of the solution and that proline insertion variants show a slight increase in phase separation as measured by a small decrease in saturation concentration (∼10% difference between wild-type and 16P at physiologically relevant salt concentrations of 75 – 150 mM), which appears correlated with the increasing length of the sequence (**Fig. 1E**). Therefore, we see no evidence for disruption of phase separation despite addition of many proline residues that should disrupt β-sheet interactions.

**Figure 1.**
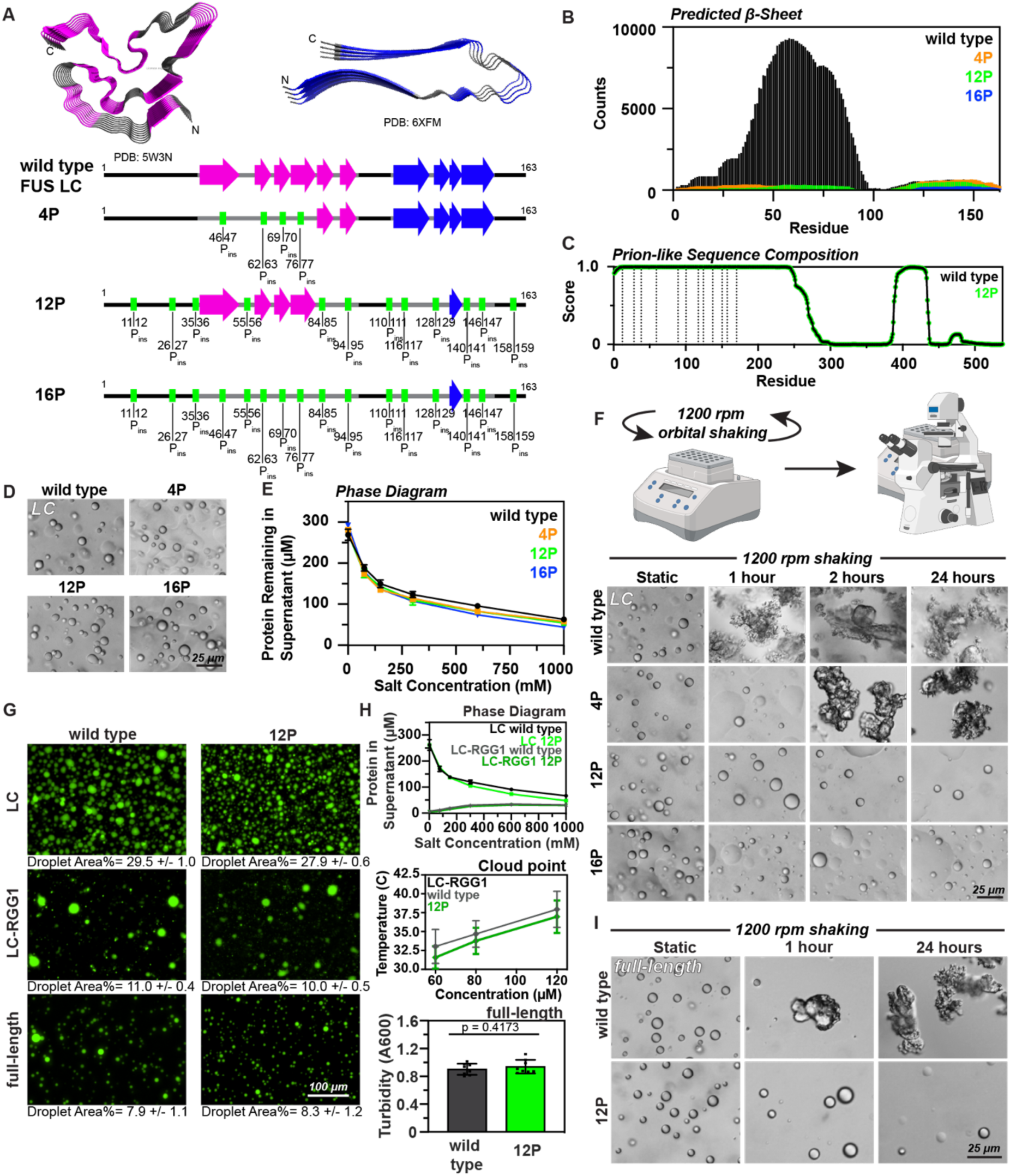
FUS aggregation, but not phase separation, is prevented by β-sheet breaking proline additions. **A)** (Top) FUS amyloid fibril core structures (5W3N and 6XFM). (Bottom) Schematic of the 4P, 12P, and 16P proline insertion mutations. Also plotted are the coincidence of the mutations with the β-sheets formed by PDB structures 5W3N and 6XFM. β-sheets determined by DSSP algorithm are represented as wide arrows, with β-strands expected to be disrupted shown by green boxes. **B)** Predicted occurrence of β-sheet structures for each residue position in FUS LC variants obtained using the β-Serpentine algorithm. Counts are generated by accumulation of the residue involvement in predicted β-sheet structures. **C)** Similarity to yeast prion sequences is not altered by polyproline insertion mutation determined using the PLAAC algorithm. A score of 1.0 signifies high similarity to the sequence composition of yeast prions. Dashed lines indicate the sequence positions where the proline insertions are made. **D)** DIC micrographs of 300 μM FUS LC variants in 20 mM HEPES, pH 7.0 with 150 mM sodium chloride. Scale bars represent 25 μm. **E)** Measurement of protein concentration remaining in the supernatant following sedimentation of 300 μM FUS LC variants in 20 mM HEPES, pH 7.0 solutions with increasing sodium chloride concentrations. Data are plotted as the mean +/- 1 s.d. using n = 3 technical replicates and confirmed with N = 2 biological replicates. **F)** (Top) schematic depicting experimental procedure. (Bottom) DIC micrographs of independent samples of 300 μM FUS LC variants under increasing durations of 1200 rpm orbital shaking at room temperature in 20 mM HEPES, pH 7.0, 150 mM sodium chloride. Scale bars represent 25 μm. **G)** Fluorescence microscopy of FUS wild type or 12P in FUS LC, FUS LC-RGG1, or full-length FUS contexts. Percentage area of droplets for each sample is quantified using n = 3 technical replicate images and verified by repeating with N = 2 biological replicates. Scale bars represent 25 μm. **H)** Phase separation of different FUS domains with and without the 12P mutation is quantified. For salt-dependent phase boundaries of FUS LC and LC-RGG1, the mean +/- 1 s.d. of n = 3 technical replicates are reported and N = 2 biological replicates were performed. For temperature dependent phase boundary of LC-RGG1, the mean +/- 1 s.d. were determined from N = 3 biological replicates consisting of n = 8 technical replicates each. For quantification of the extent of phase separation for full-length FUS variants in 20 mM HEPES, pH 7.0 150 mM sodium chloride, the mean +/- 1 s.d. were determined using n = 8 technical replicates and confirmed using N = 2 biological replicates and a student’s t-test was performed. **I)** DIC micrographs of the full-length FUS wild-type or 12P were exposed to increasing durations of 1200 rpm orbital shaking. Samples were generated as 60 μM FUS in 20 mM HEPES, pH 7.0, 150 mM sodium chloride. TEV protease was added 30 minutes prior to the experiment in order to cleave the solubilizing MBP-tag. Scale bars represent 25 μm.

To test our hypothesis that the aggregation properties of FUS could be disrupted without altering phase separation, we subjected the FUS LC proline variants to aggregating conditions using orbital agitation at 1200 rpm in a tabletop thermomixer, which causes wild type FUS LC to aggregate by 1 hr (**Fig. 1F**). Though each variant forms liquid-like droplets before shaking, aggregation as a function of shaking time of each variant was discouraged with increasing proline content. While the 4P variant formed irregularly shaped aggregates after 2 hours at these conditions, the 12P and 16P variants did not form aggregates even after 24 hours, suggesting that aggregation and phase separation could indeed be decoupled from one another. Surprised by this remarkable stability, we subjected these samples to the 1200 rpm shaking condition for 1 week and observed that the 12P and 16P samples neither formed aggregates nor ThT fluorescent assemblies over the duration of one week (**Fig S1B**). Therefore, we chose to further study the 12P variant as it showed the greatest effect with minimal perturbation to the endogenous sequence. In order to determine if this aggregation resistance could be conferred to larger truncations of FUS, we replaced the wild-type LC domain of FUS with our engineered 12P LC domain in FUS LC-RGG1, a highly phase separating segment (residues 1-284) as well as in the full-length FUS (**Fig. 1G**). In each of these constructs, liquid-like droplets formed initially were visually and quantitatively indistinguishable from the wild-type, based on analysis of the total area fraction of fluorescence microscopy images (LC: p = 0.0827; LC-RGG1: p = 0.0487; full-length: p = 0.7063). To further compare the phase separation of the wild-type and the 12P in the different domain contexts, we measured the saturation concentration of FUS LC as above as well as FUS LC-RGG1 in response to increasing concentrations of salt (**Fig 1H, top**). The matched LC and LC-RGG1 variants all phase separated nearly identically to the wild-type for the LC and LC-RGG1 variants, suggesting that the RGG1 interactions with the LC that contribute to phase separation^14^ are not altered by addition of proline in the LC. Furthermore, the inverted salt-dependent phase behavior between the LC and LC-RGG1 domain constructs that arises from charged residue interactions^14^ was similarly unaffected, suggesting that the interactions and role of the RGG1 domains were not disrupted by the presence of the proline variants in the LC. Given that proline residues can change chain expansion^63^ and are common in elastin-like and resilin-like peptides that phase separate more at higher temperatures^64^, opposite to what has been observed for FUS, we tested whether other stimuli known to affect phase separation (temperature, crowding agents) were perturbed by proline addition. To that end, we measured the cloud point of the LC-RGG1 12P variant (**Fig 1H, middle**) and visualized the phase separation of FUS LC-RGG1 in response to addition of 5% w/v polyethylene glycol (PEG) (**Fig S1C**). Similar to the result seen for increasing salt concentrations, the LC-RGG1 wild-type and 12P variant exhibited highly similar phase separation response to each of these stimuli. Finally, to determine whether the phase behavior in the full-length sequence was altered, we compared the turbidity of the full-length FUS sequence with the wild-type or 12P LC domains and found that the phase separation of the full-length protein was the same regardless of the identity of the LC domain (**Fig. 1H, bottom**). We then tested whether the 12P additions conferred aggregation resistance to the full-length FUS variant in biochemical assays. By exposing intact FUS variants to the same aggregating conditions of 1200 rpm (**Fig. 1I**), we demonstrated that full-length FUS bearing the 12P additions in the LC domain could withstand aggregation for at least 24 hours under extreme aggregation-promoting conditions (rapid shaking). This finding demonstrates that the LC domain of FUS is both necessary and sufficient to cause aggregation, and that selective modification of the primary sequence allows us to prevent aggregation while phase separation remains intact.

### β-sheet structure does not contribute to structure or motions of FUS LC in dispersed or condensed phases

Given the effects of proline on chain conformation^63^, we next tested whether the introduction of proline residues significantly altered the secondary structure or backbone dynamics of the FUS LC domain using solution-state NMR. We hypothesized that if β-sheet-forming regions contribute to the structure and motions of the FUS LC domain in the dispersed phase, then disrupting these regions with proline insertions would introduce detectable changes in structure and motions. We first examined the global protein structural fingerprint by collecting ^1^H-^15^N HSQC spectra to examine the amide ^1^H chemical shift dispersion at low concentrations where the protein is not phase separated (**Fig. 2A**). Apart from localized chemical shift perturbations near the proline insertion sites, the overall backbone amide ^1^H chemical shift dispersion remained consistent with that of a disordered region, which was also supported by effectively identical circular dichroism spectra showing random coil profiles for both sequences in the dispersed phase (**Fig S1D**). The values of the ^15^N NMR spin relaxation parameters (backbone *R*_1_ and *R*_2_ relaxation rate constants and the ^1^H-^15^N heteronuclear NOE) for the dispersed and condensed phase were measured to assess residue-specific backbone motions on the nanosecond timescale. The per-residue values were nearly identical between the wild-type and 12P, with only slightly enhanced *R*_2_ relaxation rates observed adjacent to the proline addition sites reflecting local changes in backbone structure due to proline insertions (**Fig. 2B**). The similarity in relaxation profiles suggests that the proline mutations did not introduce significant structural or dynamic perturbations to the global structure of the FUS LC domain and do not markedly “rigidify” the chain. Next, to compare the hydrodynamic properties of FUS LC 12P in the dispersed phase, we employed diffusion-ordered spectroscopy (DOSY-NMR) to measure the diffusion coefficient of the wild-type and the 12P sequences. We hypothesized that if prolines altered the degree of spatial compaction due to reduced β-sheet character, this would be reflected in changes to the diffusion coefficient. While interpreting the diffusion coefficient of IDPs in terms of their radius of hydration can be complex as solvation effects may obscure direct size comparisons^65^, such measurements provide valuable insight into the extent of chain collapse driven by intramolecular interactions. Our results revealed that the diffusion coefficient of wild-type LC (72.8 +/- 1.2 μm^2^/s) was slightly faster (by about 8.8%) than that of the 12P FUS LC (66.4 +/- 2.1 μm^2^/s) (**Fig. 2C**). This difference is consistent with the 12P sequence being lengthened from 163 residues to 175 residues (a 7.4% increase) due to the insertion of 12 prolines, which likely accounts for the reduced diffusion coefficient. Finally, we also created macroscopic condensed phases of wild-type or 12P FUS LC-RGG1 and the NMR-measured diffusion of the LC-RGG1 variants in the condensed phase was also similar (**Fig. 2D**). Simulations comparing the wild type and 12P variants of FUS LC or FUS LC-RGG1 slightly more extended conformations in the 12P variants (**Fig. 2E**) due in part to the longer sequence and a rearrangement of contacts, reducing the number of mid to long-range contacts and increasing the number of short-range contacts, especially at and near the proline insertion sites (**Fig. S2**). These simulations may also help explain why both LC-RGG1 and LC show slight chain expansion (**Fig. 2C-F**) but only FUS LC 12P shows slight enhancement in phase separation compared to the wild type sequence (**Fig. 1H**) – in FUS LC-RGG1, the 12P addition causes some disruption of interdomain (i.e. LC to RGG1) contacts that are critical drivers of phase separation^14,20^. Together, these results support the conclusion that prolines disrupting β-sheets do not directly alter the overall phase separation of FUS but rather suggests that minimizing the number of proline insertion mutations may be advantageous for the design of variants that have minimal quantitative differences in phase separation.

**Figure 2.**
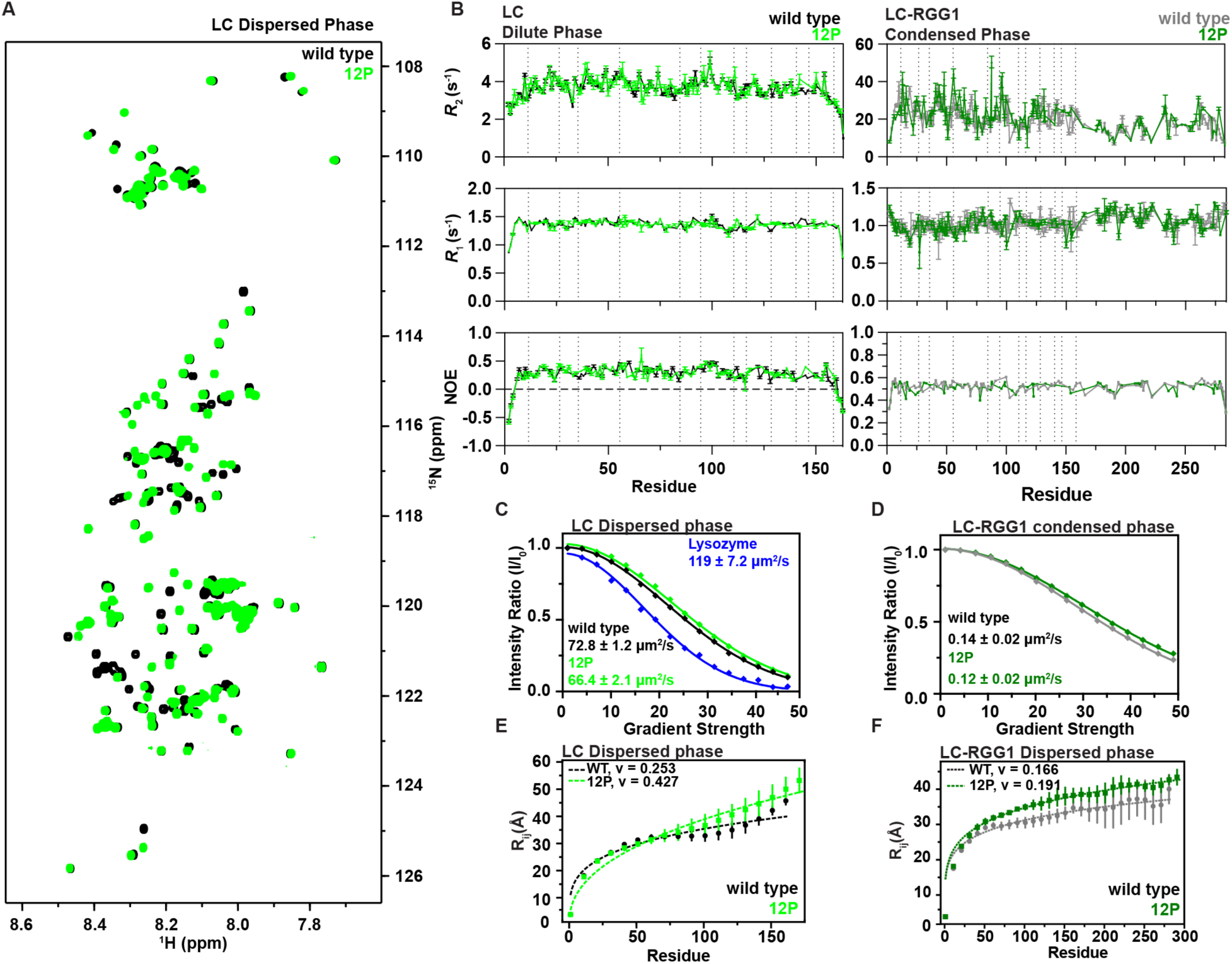
β-sheets do not contribute to global structure or motions, but slight expansion of the FUS LC backbone results from added prolines. A) Backbone amide fingerprint NMR spectra of FUS LC wild type or 12P in the dispersed phase are consistent with structural disorder and only local changes. Samples were prepared as 75 μM FUS LC in 20 mM MES, pH 5.5, 150 mM sodium chloride and 5% v/v D2O and data were collected at 298 K. B) Motions in the condensed phase measured by ^15^N spin relaxation parameters *R*1, *R2,* and heteronuclear ^1^H-^15^N NOE. Dashed lines indicate sequence positions where proline insertion mutations were made and the sequence numbering for the 12P is adjusted to match the wild-type sequence numbering to aid visual comparison. Data are plotted as mean +/- 1 s.d. C) Diffusion Ordered Spectroscopy (DOSY) NMR comparing the wild type and 12P sequences in the dispersed phase. Samples are prepared as 75 μM FUS LC in 20 mM MES, pH 5.5 with 150 mM sodium chloride and 5% (v/v) D2O lock solvent. Lysozyme is included as a biological reference. D) DOSY-NMR experiments comparing the wild type and 12P FUS LC-RGG1 diffusion coefficients in the condensed phase. E) Average intrachain distance, Rij, between the *i^th^* and *j^th^* residues computed using all-atom simulations comparing the FUS LC wild type and 12P in the dispersed phase. Standard error of the mean is computed over 3 independent trajectories for single-chain result. Dashed lines are best-fit values of R*ij* = *b* |*i – j*|*^ν^* where *ν* is the scaling exponent. F) Same as in (**E**) except comparing FUS LC-RGG1 wild-type and 12P.

### Protein backbone structure modulates aggregation and LST in FUS

Having established that prolines disrupt apparent LST in FUS without marked change to dynamics or diffusion, we sought to compare proline addition to other sequence perturbations that discourage aggregation and quantify the change in aggregation potential. Prior studies demonstrate that glycine-enriched sequences exhibit remarkable aggregation resistance^14^, for example, replacing all 42 LC serines with glycines (S→G) resulted in exceptional stability, and studies by Wang et al. show that replacing all glutamines with glycines (Q→G) similarly suppressed condensate maturation in FUS^47^. Building on these findings, we examined the aggregation behavior of the 12P variant and compared it to our S→G variant. To test this, we agitated phase separated samples of wild-type, S→G, and 12P FUS LC under orbital shaking at 1200 rpm and monitored their aggregation for 24 hours (**Fig. 3A,B**). Interestingly, although the S→G condensates maintained their liquid-like quality much longer than the wild type before transitioning into dynamically arrested condensates which appeared dense and irregularly shaped, these assemblies did not exhibit classical aggregated morphology and showed only weak fluorescence enhancement with the amyloid-detecting thioflavin T (ThT). After 24 hours, the structures formed by the S→G variant progressed to slightly higher ThT fluorescence, suggesting that large sequence changes may prevent ThT-based detection of aggregation. In contrast, the 12P condensates again showed remarkable stability, retaining their spherical morphology and liquid-like appearance even after 24 hours, with no evidence of aggregation or ThT fluorescence. These results suggest that while the backbone structure of glycine-enriched sequences provide some resistance to aggregation, selective insertion of prolines into β-sheet regions provide even greater stability by preventing aggregation-prone regions from forming cross-β structures. Furthermore, previous studies have also shown that glycine enrichment significantly decreases the saturation concentration of FUS LC by promoting interactions between residues near glycine positions^14^, which may cause unwanted effects on function, whereas our data (**Fig. 1D-H**) demonstrate that the saturation concentration of the 12P variant is minimally altered by the addition of proline.

**Figure 3.**
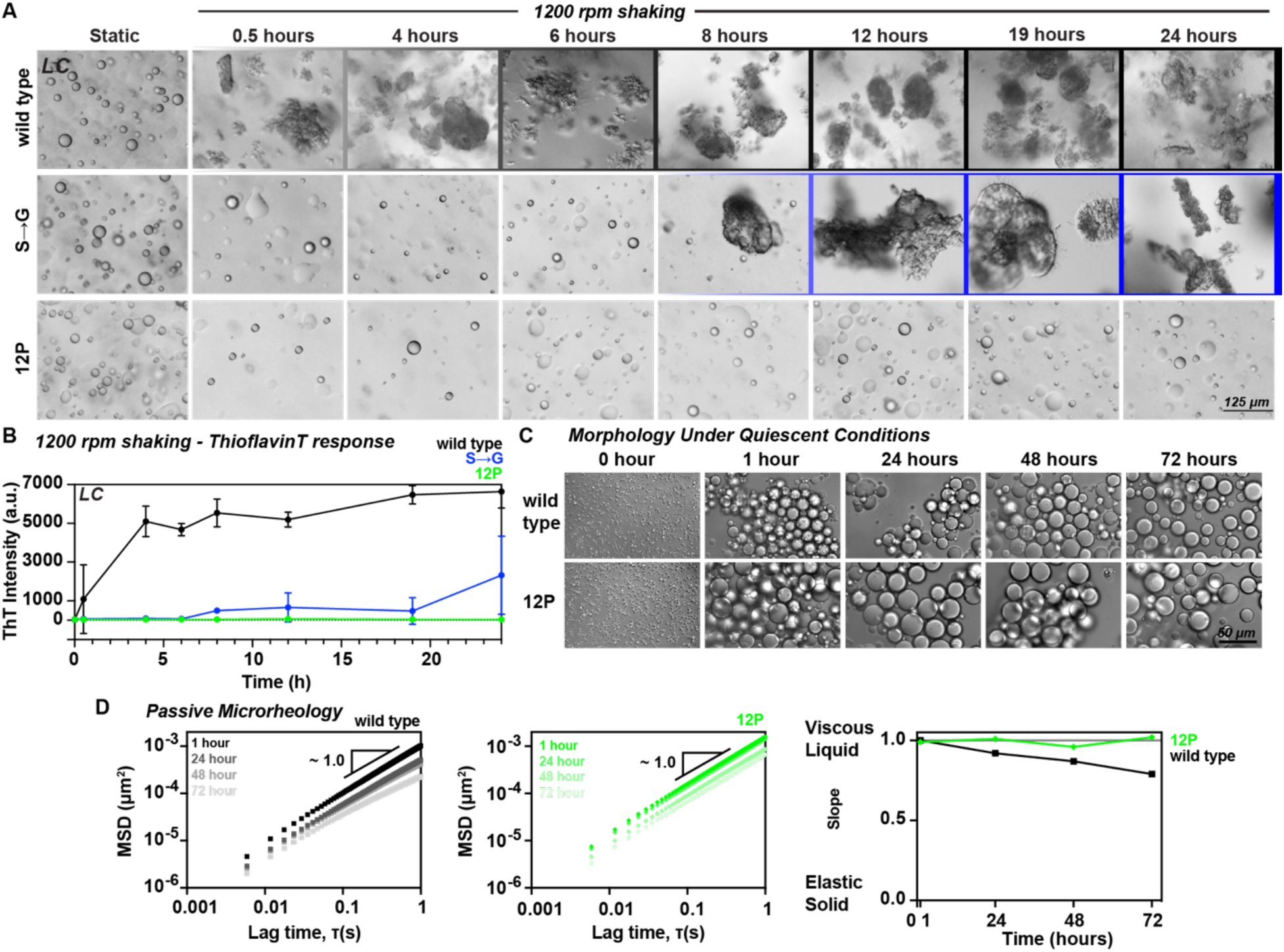
Protein backbone structure modulates aggregation and LST in FUS. **A)** Independent samples of 300 μM FUS LC wild-type, S→G, or 12P were subjected to 1200 rpm shaking for up to 24 hours to induce aggregation. DIC micrographs were taken at increasing time points using differential interference contrast (DIC) microscopy. **B)** Thioflavin T fluorescence of the FUS LC variants in (**A**) was monitored up to 24 hours for independent samples continuously agitated at 1200 rpm. Data are reported as mean +/- 1 s.d., n = 3 technical replicates. **C)** Brightfield micrographs of FUS LC-RGG1 condensates under quiescent conditions for up to 72 hours. Samples were prepared as 60 μM FUS LC-RGG1 in 20 mM HEPES, pH 7.0, 150 mM sodium chloride and transferred to a 96-well plate that was kept at room temperature over the duration of the experiment. **D)** Passive particle-tracking microrheology was used to measure the change in viscoelasticity of LC-RGG1 condensates over the course of 72 hours for the wild type (left) and 12P (center) condensed phase. The slope of the log-log plots corresponds to the condensates’ viscoelastic properties where 1.0 is for viscous liquids and 0 for elastic solids (Right).

We next sought to understand how disrupting local backbone structure impacts the LST in FUS condensates. Previous studies have demonstrated that the accumulation of persistent inter-protein β-sheets drives the aging of biomolecular condensates^43^. We hypothesized that disrupting β-sheet formation would prevent the development of these higher-order networks that facilitate the LST, thereby inhibiting hysteresis. To test this, we used video particle-tracking passive microrheology to monitor the mechanical properties of wild-type and 12P LC-RGG1 condensates over a 72-hour quiescent aging period (**Fig. 3C,D**). By optically tracking fluorescent polystyrene beads embedded in the condensates, we measured the mean-squared displacement (MSD) of the beads and computed the diffusivity exponent (α) (see Methods). In these experiments, a scale-invariable MSD over lag times with α = 1 is representative of a viscous liquid, while α < 1 is characteristic of viscoelastic behavior. Consistent with previous reports, wild-type condensates exhibited subdiffusive motion of fluorescent beads, accompanied by a corresponding decrease in slope α of MSD curves over the aging period, indicative of the formation of a more rigid network as the wild-type condensates aged (**Fig. 3D**). In contrast, 12P condensates maintained consistent mechanical properties throughout the 72-hour period, showing no signs of physical aging. These results suggest that the disruption of the local backbone structure prevents the formation of higher-order network assembly in condensates that drive condensate irreversibility and aggregation.

### Localization to nuclear foci and transcriptional regulation are not mediated by β-sheets

The diverse roles of FUS in cellular processes include the regulation of transcription and RNA processing as well as the integrated stress response^7,8^ and are well-documented. However, it remains debated whether β-sheet structures contribute to physiological FUS function or merely reflect the limited structural encoding of low-complexity sequences^66^. Studies on yeast prions, including pioneering work by Shorter and Lindquist^67^, have shown that reversible fibrillization underpins the epigenetic inheritance conferred by prions in yeast^68^. Whether similar fibrillization events or even more transient or “labile” β-sheet structures are required for the diverse functions of human FUS, or if instead β-sheet structure contributes exclusively to toxicity, remains unsolved. One well-characterized function of FUS is its ability to form biomolecular condensates that recruit the C-terminal disordered tail of RNA pol, leading to transcriptional activation^2,69^. This partitioning process depends on FUS self-interactions, as well as interactions between FUS and RNAPII, raising the question of whether β-sheets might contribute to the interactions required for assembly of transcriptionally active condensates and recognition of transcription factors like RNAPII. First, we compared the partitioning of either Alexa488-tagged FUS LC or the last 27 heptads and C-terminal tail of human RNA pol II^70^ into condensates formed by wild-type or 12P FUS LC. By measuring the fluorescence of protein remaining in the supernatant after condensate formation (**Fig. 4A**), we observed that the partitioning of both Alexa488-FUS LC (wild type) and Alexa488-RNA pol II was nearly identical (p = 0.95 for Alexa488-FUS LC; p = 0.66 for Alexa488-RNAPII) in wild-type and 12P condensates (**Fig. 4B,C**). These results suggest that β-sheet interactions are not critical for the multivalent interactions driving host-client partitioning or condensate assembly. In light of this, we next sought to test if the absence of β-sheets alters the molecular organization of FUS-containing condensates in a cellular environment. We focused on paraspeckles, ribonucleoprotein granules that regulate nuclear RNA retention, because their assembly is known to be sensitive to perturbations of the FUS LC domain^37,71,72^. To test if β-sheet formation within the FUS low-complexity domain contributes to its recruitment to paraspeckles, we created a genome-edited U2OS cell line expressing eGFP-tagged mouse 12P FUS from the endogenous locus, thus ensuring physiological levels of protein expression (**Fig. S3A-D**). Non-edited cells (NEC) and cells expressing wild-type eGFP-FUS, as described in our parallel work^37^, served as controls. Fluorescence microscopy revealed extensive co-localization of both wild-type and 12P eGFP-FUS with the paraspeckle marker PSPC1 in nuclear condensates (**Fig. 4D**). Indeed, one-dimensional fluorescence intensity profiles across U2OS nuclei showed a strong correlation between eGFP-FUS and PSPC1signals for both constructs. Furthermore, the number of eGFP-positive foci present in either wild-type or 12P cells was statistically indistinguishable (p=0.27). Together, these results indicate that the β-sheet-forming ability of FUS is dispensable for its recruitment to PSPC1-positive foci. Given that localization to these foci strictly depends on the phase separation ability of FUS as we show by tyrosine-to-alanine variant in this exact assay in our parallel work^37^, these findings further strengthen our previous conclusion that the proline insertions do not impair the partitioning of FUS into liquid-like condensates.

**Figure 4.**
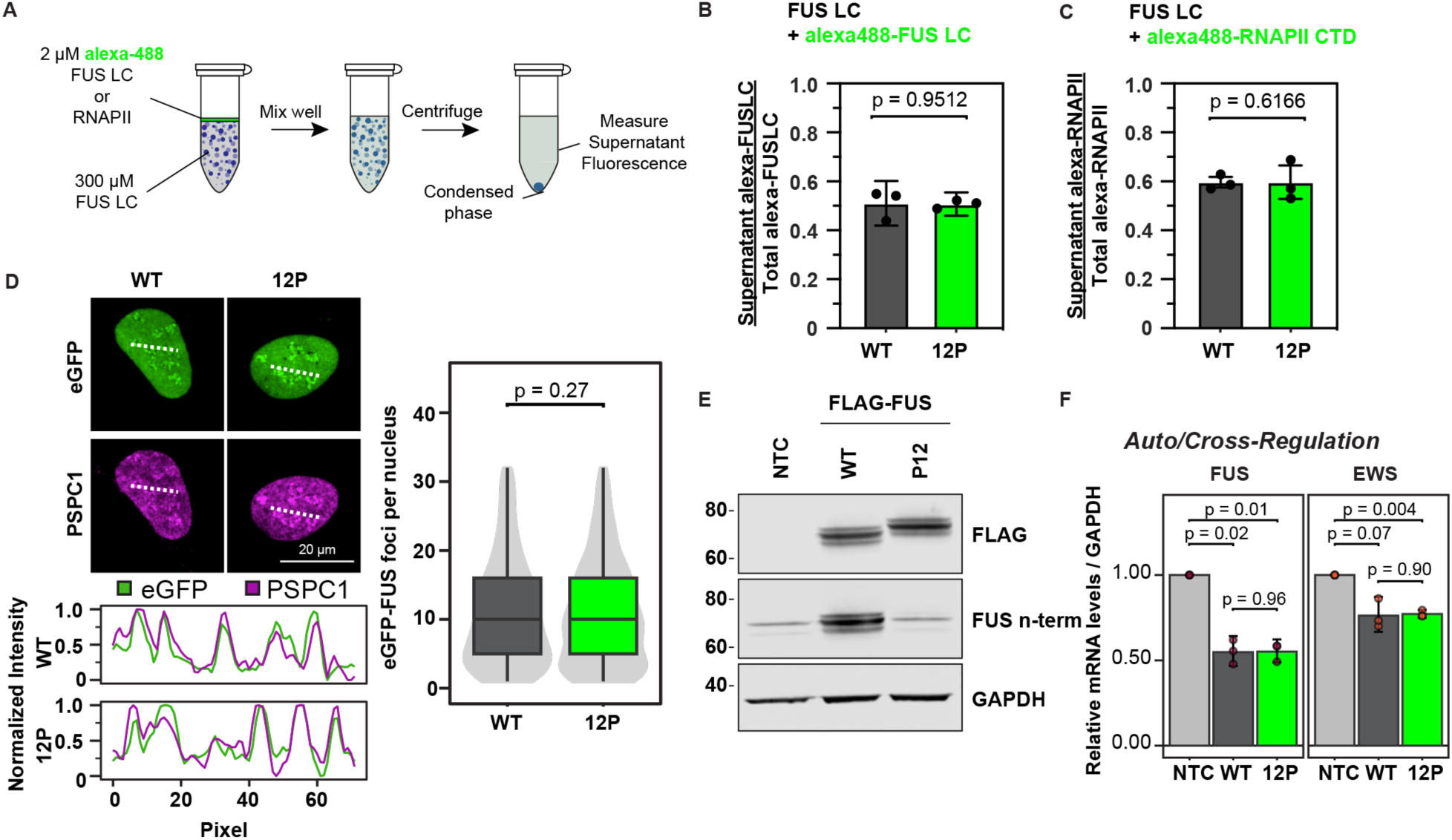
Nuclear localization and transcriptional regulation are not mediated by β-sheets. A) Schematic showing how the partitioning experiment for (**B)** and (**C)** is designed. B) Comparison of Alexa488-FUS LC self-partitioning into FUS LC wild-type or 12P condensed phase. Data reported as mean +/- 1 s.d., n = 3 technical replicates, N = 2 biological replicates. C) Comparison of Alexa488-RNA Pol II partitioning into FUS LC wild-type or 12P condensed phase. Data reported as mean +/- 1 s.d., n = 3 technical replicates, N = 2 biological replicates. D) Fluorescence microscopy images showing eGFP-FUS (green) and PSPC1 (magenta) in U2OS cells expressing wild-type or 12P eGFP-FUS. Fluorescence intensity profiles for each channel were taken along the dashed lines indicated in the images and rescaled from 0 to 1 to examine colocalization. The violin-boxplot shows the number of eGFP foci per foci-positive nucleus. The plot displays median lines, interquartile range (IQR) boxes and 1.5 × IQR whiskers. n = 523 (WT) and 493 (12P) nuclei, from N = 3 independent biological replicates. Statistical significance was determined by unpaired t-test using the mean number of eGFP-FUS foci of each biological replicate. E) Western blot analysis of FLAG-FUS expression levels with GAPDH as loading control. Anti-FLAG antibodies detect total FLAG-FUS, while an N-terminal FUS antibody verifies the presence of proline insertions that disrupt epitope recognition. F) Endogenous FUS and EWS mRNA levels in HeLa cells expressing FLAG-tagged WT or 12P FUS determined by RT-qPCR relative to non-transfected control cells (NTC) to compare auto and cross-regulation activity. Data are reported as mean +/- 1 s.d. from N = 3 independent biological replicates. P values were calculated with ΔΔCt values by one-way ANOVA followed by Tukey’s Honest Significant Difference (HSD) test for multiple comparison

Next, we investigated the role of β-sheets in FUS function in gene expression, focusing on two physiologically relevant events: (1) FUS autoregulation, which occurs through retention of introns 6 and 7^73^ and depends on LC-mediated FUS phase separation^1^; and (2) cross-regulation of EWS, a paralog of FUS and member of the FET family (FUS, EWS, and TAF-15), through a mechanism that remains poorly understood. To test whether these regulatory RNA processing events rely on β-sheet formation, we overexpressed FLAG-tagged wild-type and 12P FUS in HeLa cells, achieving similar expression levels (**Fig. S3C**), and measured endogenous FUS and EWS mRNA levels by RT-qPCR. Both mRNAs were significantly downregulated upon overexpression of wild-type and 12P FUS (**Fig. 4F**), indicating that β-sheets do not play an integral role in these regulatory events. Importantly, autoregulation is impaired by disruption of the phase separation potential of FUS LC by tyrosine-to-serine substitutions^1^, highlighting the importance of the LC domain but not its β-sheet propensity for RNA processing functions.

### Ability to form β-sheets is dispensable for FUS function in DNA damage response

Besides its various roles in RNA metabolism, FUS also participates in the DNA damage repair pathways for both single-strand (SSB) and double-strand (DSB) breaks^10^. Robust accumulation of FUS at sites of DNA damage requires the LC domain^74^ and its phase separation ability^37^ as shown in our parallel experiments. Hence, we first assessed the impact of reducing β-sheet content in FUS on its recruitment to DNA damage sites. Here, we subjected the U2OS cell lines to laser microirradiation and monitored eGFP-FUS localization by live-cell imaging. Both wild-type and 12P FUS were rapidly recruited to damage sites, demonstrating similar dynamics and phase transition behavior in response to DNA breaks (**Fig. 5A**).

**Figure 5.**
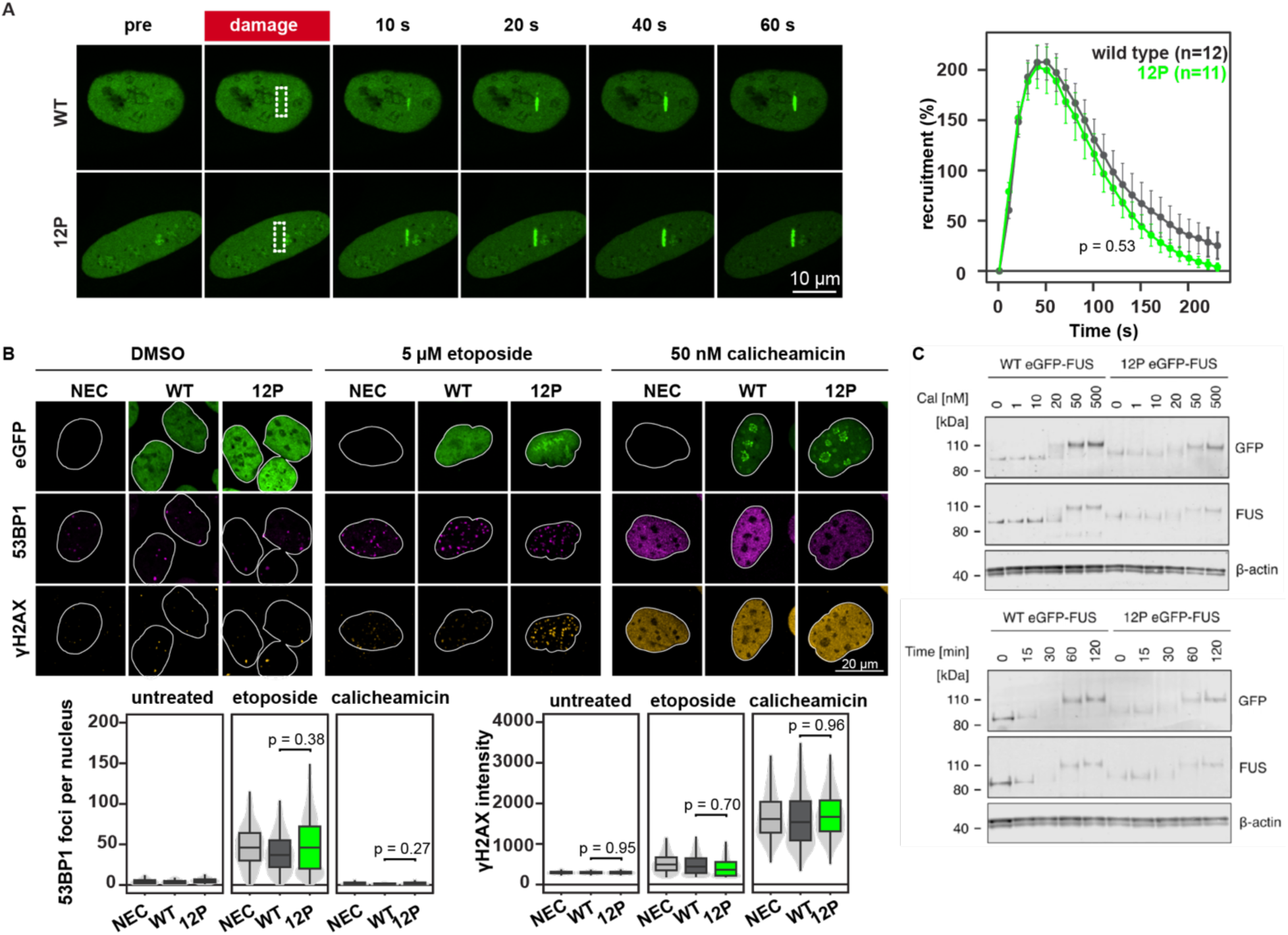
DNA damage response and calicheamicin-dependent FUS phosphorylation are not altered by β-sheets. **A)** U2OS cells expressing eGFP-FUS wild-type or 12P were used to compare the localization of FUS to DNA strand break sites induced by excimer laser irradiation (left). Images were recorded up to 240 seconds and the total recruitment was quantified at 10-second intervals (right). Scale bar, 10 µm. Data are reported as mean +/- SEM of n = 12 (WT) and n = 11 (12P) nuclei, taken from N = 3 independent experiments. For statistical analysis, recruitment was quantified as area under the curve, and differences between wild-type and P12 nuclei were assessed using a linear mixed-effects model with biological replicate as a random effect. **B)** Fluorescence microscopy images showing the localization of eGFP-FUS (green), 53BP1 (magenta) and γH2AX (orange) in DMSO-treated control cells (left) and upon induction of DNA damage with etoposide (center) or calicheamicin (right). Scale bar, 20 µm. Violin-boxplots show number of 53BP1 foci per positive nucleus. The plot displays median lines, interquartile range (IQR) boxes and 1.5 × IQR whiskers. DMSO-treated: n = 3,566 (NEC), 2,783 (WT), 3,226 (12P); etoposide-treated: n = 1,767 (NEC), 1,804 (WT), 2,178 (12P); calicheamicin-treated: n = 451 (NEC), 447 (WT), 475 (12P) nuclei, from N = 3 independent biological replicates. Statistical tests were performed by one-way ANOVA followed by Tukey’s Honest Significant Difference (HSD) test for multiple comparison. **C)** Western blot analysis of eGFP-FUS following 1h exposure to increasing concentrations of calicheamicin for wild-type FUS and the 12P variant (top). The time required to observe FUS phosphorylation after exposure to 50 nM calicheamicin was compared for FUS wild-type and 12P (bottom). β-actin served as loading control.

To further evaluate whether β-sheet formation affects the DSB response in our system, we used automated high-content imaging to examine serine 139-phosphorylated histone H2AX (γH2AX), a canonical marker of early DNA damage signaling, and 53BP1, a downstream response mediator that promotes the DSB non-homologous end joining (NHEJ) pathway. In untreated cells, we observed no difference in γH2AX levels and the number of 53BP1 foci between wild-type and 12P FUS-expressing cells, as well as non-edited controls (NEC) (**Fig. 5B**) Given that FUS depletion^10^ or selective inhibition of FUS phase separation^37^ are linked to increased γH2AX levels, this result suggests that β-sheet structures do not play an important role in this process. Consistently, upon induction of DNA damage with the topoisomerase inhibitor etoposide or the DNA-cleaving antibiotic calicheamicin, we observed the expected increases in γH2AX levels, but again no differences between wild-type and 12P FUS cells (**Fig. 5B**). Likewise, the number of 53BP1 foci was strongly increased in response to etoposide, independently of the genotype of the cells (**Fig. 5B**). While calicheamicin treatment did not strongly promote 53BP1 foci formation, we noted that it induced a marked redistribution of wild-type and 12P eGFP-FUS to perinucleolar condensates, a behavior that relies on a functional FUS LC domain and phase separation ability^37^. In addition to inducing this re-localization of FUS within the nucleus, calicheamicin treatment is also known to induce phosphorylation of the FUS LC domain via DNA-dependent protein kinase (DNA-PK)^75^. To assess if the 12P additions affect this post-translational modification, we subjected our U2OS cells to increasing concentrations of calicheamicin. Using Western blots, we observed no detectable differences in eGFP-FUS phosphorylation of 12P relative to wild-type, as indicated by large shifts in electrophoretic mobility (**Fig. 5C**). Similarly, exposure to a fixed amount of calicheamicin yielded no apparent differences in the kinetics of eGFP-FUS phosphorylation (**Fig. 5C**).

To probe how the aggregation-deficient variant affects condensate maturation and FUS dysfunction, we attempted to induce mature stress granules in U2OS cells using poly(I:C) as others have reported^76^, but failed to form persistent granules even for wild-type FUS, likely due to cell-type differences (**Fig. S5,6**). A modified protocol using heat-shock followed by a low dose of sodium arsenite produced persistent assemblies with only a modest increase in granule reversibility for FUS 12P compared to wild type, underscoring that cell line and poly(I:C) properties, concentration and treatment duration, strongly influence different responses to viral nucleic acid mimics.

In summary, our experiments indicate that β-sheet formation does not contribute to FUS localization, post-translational modification, and function in the DNA damage response.

### Inhibiting β-sheets entirely rescues toxicity in Drosophila models

Studies of FUS toxicity in *Drosophila* models have shown that mutant FUS variants promoting cytoplasmic accumulation cause various functional defects, including retinal degeneration, locomotor dysfunction, and lethality^35^. Some studies propose that compensatory switching between the ubiquitin proteasome system (UPS) and autophagy becomes dysregulated following altered interaction between NLS-mutant FUS and HDAC6^77^, while others suggest that activation of the unfolded protein response (UPR) pathway drives mitochondrial toxicity^78^. However, it is unclear if formation of cytoplasmic aggregates is required for toxicity or if mislocalization alone is sufficient to drive toxicity, as has been suggested for TDP-43^79^. To identify the link between cytoplasmic localization, aggregation, and toxicity, we engineered human FUS 12P either without or with an additional P525L mutation that prevents proper nuclear localization (wild-type, 12P, P525L, and P525L/12P double mutant) in a *Drosophila* model driving expression in motor neurons and the eyes using tissue-specific drivers.

First, we examined the effects of β-sheet propensity on development for exogenous FUS-expressing animals^80–82^. When the wild-type or P525L variants were expressed in the eyes, the degenerated ommatidial phenotype was observed (**Fig. 6A, S4B**) as observed previously for these variants, whereas the variants expressing the 12P or the P525L/12P FUS completely matched the non-expressing control. We then used flies expressing FUS with a motor neuron driver. When examining the wing structure of the surviving flies, a dramatic distortion of wing development (a “curly-wing” phenotype) was identified for the wild-type and P525L flies. When flies expressed either the 12P variant or the P525L/12P double variant, this effect was entirely rescued (**Fig. 6B**). Next, we examined pupal eclosion of these FUS-expressing animals. Flies expressing the wild-type or P525L FUS showed strong defects in pupal eclosion compared to the control that were dramatically reversed by incorporation of the 12P modification, showing a complete rescue to the non-FUS-expressing control levels (**Fig. 6C**). We performed Western Blot analysis of our fly lines to compare the FUS expression levels (**Fig. 4E**) and found that the variants showed similar levels with the exception of the P525L/12P double variant which had a higher expression level than other variants. Interestingly, despite the higher expression, the P525L/12P variant did not show eclosion defects that are prominent in FUS P525L expressing animals, suggesting that prevention of β-sheet formation dramatically reduces toxicity.

**Figure 6.**
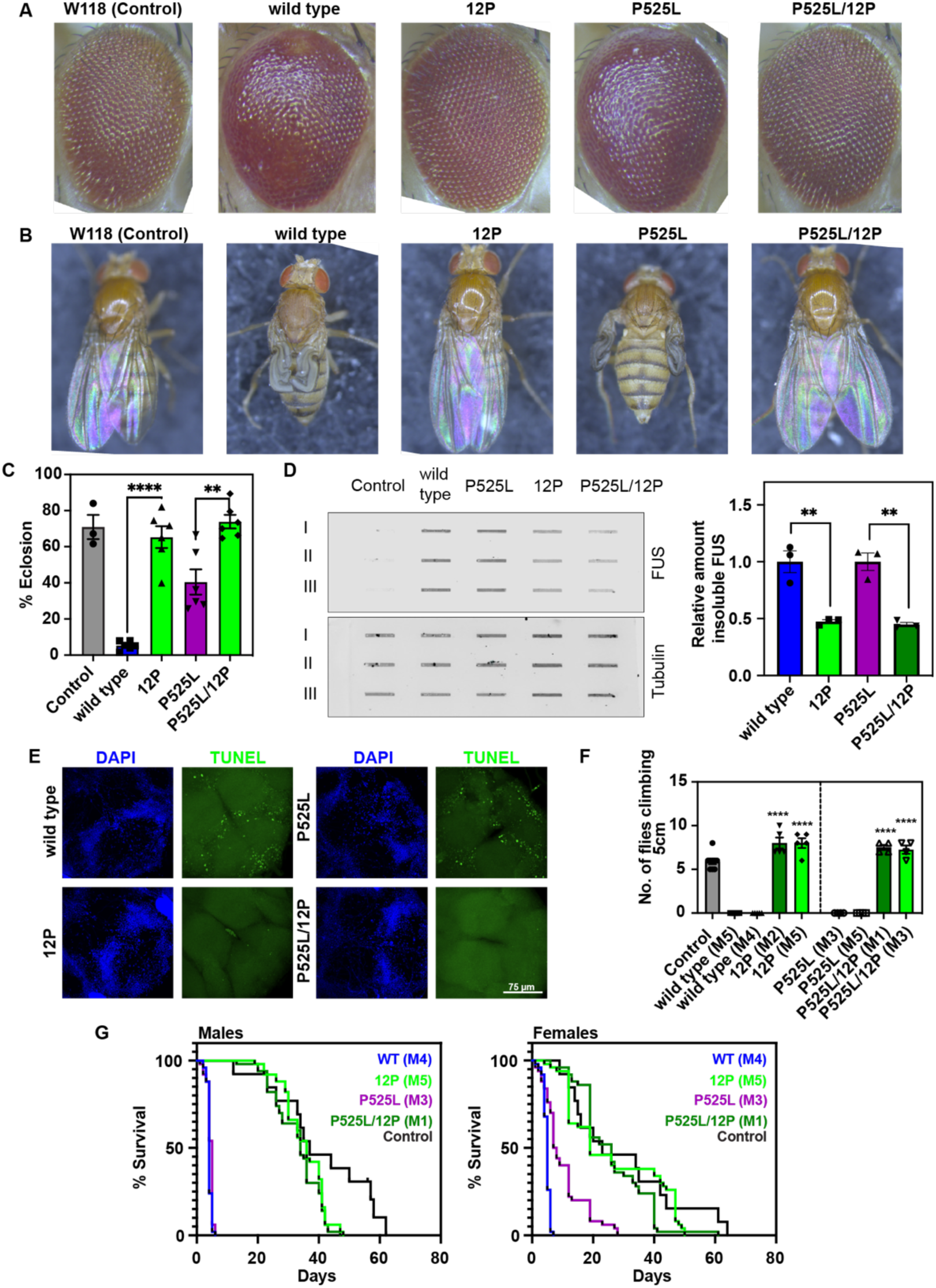
Inhibiting β-sheet formation entirely rescues FUS-induced motor dysfunction and reduced lifespan in *Drosophila* models. **A)** Images of *Drosophila* eyes for flies expressing FUS variants using an eye-specific driver (GMR-gal4 driver). **B)** Images of the wing development of *Drosophila* expressing wild-type, 12P or P525L, P525L/12P through a motor neuron driver (D42-gal4). **C)** Percent eclosion of *Drosophila* larvae for flies expressing a control or FUS wild-type, 12P, P525L, or P525L/12P with a GMR-Gal4 motor neuron-specific driver (D42-gal4). Data are plotted as mean +/- standard error of the mean using n = 6 technical replicates and compared for statistical significance using a student’s t-test (**** represents P < 0.0001 and ** represents P <0.05). **D)** (Left) Cellulose acetate filter-trap of total cell fly head lysate immunostained for FUS to visualize insoluble assemblies. (Right) Relative quantification of filter trap band intensities. Comparisons were made using an unpaired t-test (** represents P <0.05). **E)** TUNEL and DAPI staining of Drosophila brain cross-sections for flies expressing FUS variants using a motor neuron-specific driver. Scale bar represents 75 μm. **F)** The climbing ability of the flies were compared by measuring the ability to climb 5 cm for flies expressing FUS variants through a GMR-Gal4 motor neuron-specific driver. Data are plotted as mean +/- standard error of the mean. Statistical significances between groups were compared using Šídák’s multiple comparisons test. **** represents P < 0.0001. **G)** Comparison of lifespan for male and female flies expressing FUS variants.

To determine if reduced toxicity associated with 12P is linked to reduced aggregation, we performed fractionation experiments by passing total fly head lysate through cellulose acetate filters to measure the amount of solid assemblies containing FUS (**Fig. 6D**). We observed about half of the amount of FUS retained in the filter membrane for the 12P and P525L/12P variants compared to the wild-type or P525L variants. Dissections of the fly brains were stained for dead neurons using TUNEL, and interestingly, the brains of the flies expressing the 12P variant did not show any obvious signs of neuronal death (**Fig. 6E**). We interpret these findings to suggest that more aggregated FUS was retained for the wild-type or P525L compared to the 12P variants. In this case, the overall lower amount of 12P FUS variants retained in the filter suggests that there is less solid FUS being formed, which may explain the decrease in toxic phenotypes observed in this model system.

To examine the impact of FUS aggregation on motor neuron degeneration, these FUS variants were expressed in motor neurons and the flies were subjected to a previously reported motor function climbing assay^81–83^. In this experiment, we observed the number of flies that were able to reach 5 or 10 cm (**Fig. 6F** and **S4C**, respectively). We observed that the flies expressing the FUS wild-type or P525L background moved very little, whereas the flies expressing either 12P or the P525L/12P climbed significantly better compared to the FUS P525L animals (**Fig. S4C**). These data suggest that expression of wild-type or P525L FUS in these systems is highly toxic; however, this toxicity is drastically reduced in the 12P variants.

Given that FUS mutations lead to aggressive and juvenile onset ALS, we examined the impact of 12P with and without P525L on the fly lifespan. Flies expressing the wild-type or P525L FUS had an average lifespan of about 5 – 10 days, except for the female flies expressing the P525L variant which survived for about 20 days, suggesting a sex-specific effect for FUS P525L toxicity (**Fig. 6G**). This sex-specific difference may result from differential expression of the Xrp1 gene, which has been shown to be positively correlated with FUS orthologue Caz toxicity, and its subsequent comparatively enhanced expression in the male *Drosophila*^84^ due to dosage compensation. Regardless, flies expressing the 12P modification in either case showed a significant increase in lifespan and were nearly indistinguishable from the non-expressing controls. Together, these results show that removing the ability to form β-sheets from the FUS LC sequence prevents FUS cellular and organismal toxicity in fly models.

## Discussion

Biomolecular condensates have been increasingly recognized for their roles in critical cellular functions but also their potential contribution of the condensates or their RNA-binding protein components to pathology, particularly in neurodegenerative diseases associated with protein aggregation. Therefore, understanding the molecular mechanisms governing the transition from dynamic liquid-like states to solid-like fibrillar inclusions, how they contribute to function, and the determinants of reversible assembly are essential to characterize. The current molecular views for phase separation and aggregation of proteins with prion-like domains focus on molecular grammars and solved structures of fibrils cores, respectively, though not much is known about how these states interconvert. Some frameworks posit that the β-sheets that are found in aggregates are structurally related to local β-sheet structure in the form of steric zippers and LARKS that can transiently form to stabilize phase separation. Competing frameworks suggest β-sheet structure is absent in the dynamic liquid-like forms but accumulate in the liquid to solid transition, which may nucleate at the liquid-liquid interface of the condensate. Additionally, precisely how FUS exerts gain-of-function toxicity is widely disputed, some theories suggest that the aggregates themselves are cytotoxic, while other viewpoints suggest that mislocalization alone is sufficient for toxicity. Here, we set out to determine the link between β-sheets and the LST, and how the LST drives function and toxicity in cellular and organismal models.

First, we employed a structure-breaking design approach to engineer a FUS variant adding proline residues in sequence regions predicted to form β-sheet structures. We found that proline mutations in the LC domain of FUS prevent aggregation of FUS but do not change the disordered global structure or motions in the dispersed phase, and do not perturb the phase behavior in the LC alone or when LC domain of the full-length FUS protein is replaced by our engineered LC domain. These data conclusively support the view that the LC domain of FUS is truly disordered both as a monomer and in the condensed phase, even in regions predicted to have a high propensity to form β-sheet structures, and that the formation of these motifs is not important for the phase behavior of FUS. This is in contrast to previous indirect studies suggesting that reversible amyloid structures enriched in β-sheet are responsible for stabilizing phase separation^85,86^, but supports other findings on phase separation suggesting that dynamic multivalent interactions are the dominant forces governing phase separation^15,87,88^.

Second, our experiments in test tubes show that our engineered variant is highly resistant to extreme aggregation-promoting conditions. The approximately uniform distribution of prolines throughout the FUS LC sequence appeared to confer these anti-aggregation properties when compared with the 4P variant that only targeted one prominent amyloid core in the LC domain, likely due the formation of new core regions or rearrangement of existing cores that were able to form when shaking the 4P variant. This alteration in stability against aggregation that we observed is based on structural changes to the protein backbone. Indeed, FUS LC sequences enriched in glycine exhibited strong aggregation-delaying properties, even more so than the 4P. Interestingly the total ThT intensity of the structures formed by the S→G variant appeared to lag behind the visual observation of non-spherical droplets that lacked liquid-like qualities, suggesting that loss of liquidity happened prior to formation of robustly ThT-positive assemblies and hence that ThT fluorescence does not correlate perfectly with “aggregation” for glycine-rich sequences. Addition of prolines also preserved the liquid-like microrheology over long times, suggesting that condensate maturation likely results from the accumulation of stable β-sheet cross-links that slowly accumulate over time. We interpret the absence of this microrheological change in the 12P variant to be due to the disrupted ability to form β-sheet structure that drive persistent intermolecular association. Importantly, we show that these aggregation-resistant mutations in the LC domain can confer protective effects to the full-length FUS sequence, demonstrating that preventing aggregation of the LC domain for FUS is sufficient for preventing aggregation of the entire FUS protein and hence aggregation of FUS requires the LC domain’s β-sheet forming ability. The relationship of proline to reduced aggregation has also been studied in amylin where the primary sequence of mouse and rat amylin is resistant to amyloid formation compared to the human form of amylin^56,57^, in good agreement with these studies.

Third, when we tested the cellular functions of FUS containing our engineered LC domain, we found that our variant had no effect on basic cellular functions of FUS indicating that the β-sheet regions predicted to form in the aggregated state of FUS did not play a functional role in cell models. In particular, phase-transition-dependent recruitment of FUS to nuclear compartments, like paraspeckles, and to sites of DNA damage showed no biologically significant differences. This similarity is underscored by the capacity of FUS to regulate its own expression, as well as that of EWS, another member of the FET family of proteins. This is in contrast to other studies suggesting that β-sheets are required for functional phase separation of other prion-like RNA-binding proteins^24,44^ and even long-term memory in *Drosophila*^89^ and *Aplysia*^90^ models. This further supports the view that non-liquid states along the LST formed by β-sheets are not necessary for FUS cellular functions, which challenges claims that suggest that gelation and viscoelastic transitions directly play a role in FUS function^43^. However, decreased diffusion and loss of liquidity will certainly reduce the movement and exchange of guest molecules within the phase, which may have context-specific effects on condensate function.

Fourth, we found striking differences in the toxicity profile of FUS when our engineered LC domain was substituted for the endogenous FUS LC domain. For flies expressing either the wild-type or a disease-linked P525L FUS background, presence of the 12P mutation produced a complete rescue of the toxic phenotype in all of our fly model systems. This was accompanied by enhanced survival in a fly model, even for female flies, which showed unexpected longer survival for FUS P525L variant compared to FUS wild-type. These findings show that the toxicity of FUS is directly linked to the formation of solid aggregates that are β-sheet rich in nature and that FUS localization to the cytoplasm alone does not exert significant toxicity. Upon inspection of fly brain cross-sections, we find an absence of dead neurons for the engineered 12P variant and show that the FUS 12P and P525L/12P dramatically reduce the cellular burden of solid inclusions (despite comparable or even enhanced total protein levels) associated with toxicity. These findings support the idea that the emergence of β-sheet structures can activate cellular stress response pathways that ultimately drive gain-of-function toxicity.

Current clinical trials treating FUS-ALS employ the use of antisense oligonucleotides (ION363) to knock down FUS in juvenile patients carrying the cytoplasmic localizing P525L mutation^91^. These clinical trials thus far have demonstrated the potential for remarkable effectiveness and disease slowing, but because of the experimental nature of this treatment modality and the limited number of in-human studies, no longevity studies have been performed to date. Because other lab-based cellular studies have shown that FUS knockdown leads to cell cycle arrest due to its well-characterized function as a regulator of transcription^92^, it is uncertain what the outcomes of these patients will be if they are required to endure life-long FUS silencing. Hence, administering a genetic or RNA therapy to express a functional but aggregation-resistant variant like the FUS 12P could effectively serve as a FUS replacement therapy and could be supplemented between FUS ASO treatments as a therapeutic strategy to maintain FUS-dependent cellular functions. Importantly, therapies designed to express the aggregating-proof but fully functional variant could be a viable strategy on its own due to downregulation of aggregation-prone (wild-type and ALS-associated variants) via RNA-splicing dependent autoregulation. Finally, these data suggest that therapeutics targeting defined β-sheet structures, even without relocalization to the nucleus, could offer significant benefit, as they may allow specific depletion of toxic species without loss of functional ones.

Together, these findings show a robust decoupling of phase separation and aggregation in FUS and demonstrate a novel mechanism through which function and toxicity of FUS can be probed. These data demonstrate that proline addition is an effective method to study functional aspects of proteins with prion-like domains that are exceptionally difficult to characterize due to their aggregation-prone nature. Finally, these findings show a path to treat that diseases caused by proteins that are both strongly aggregation-prone and whose protein levels are strongly autoregulatory, such as TDP-43, by exogenous expression of an aggregation-proof variant.

## Acknowledgements

We thank Dr. Mandar Naik for NMR assistance and the Structural Biology Core Facility at Brown University. We also thank George Chennel and Chen Liang from the Wohl Cellular Imaging Centre (King’s College London, London, UK) for training and technical support with all the microscopy techniques used for cell-based assays. This work was supported in part by the National Institute of General Medical Sciences R01GM147677 (to N.L.F.) and R35GM153388 (to J.M.). N.W. and R.P. were supported in part by an NIGMS training grant (T32GM139793). N.W. was supported in part by a Blavatnik Family Fellowship from Brown University. J.A. was supported by funding awarded from the Motor Neurone Disease Association (grant 911-792, to MD.R). D.J. was supported by an Alzheimer’s Research UK fellowship (ARUK-RF2024-002). This work was also made possible through the support of the NOMIS foundation (to MD.R) and the UK Dementia Research Institute (grant UKDRI-6204, to MD.R), through UK DRI Ltd, principally funded by the Medical Research Council. We also thank the Texas A&M High Performance Research Computing (HPRC) for providing essential computational resources.

## Author Contributions

N.W. and N.L.F. designed the 12P sequence and conceptualized the study. N.W designed and performed the biochemical analyses and wrote the manuscript with guidance and feedback from N.L.F. D.A.B. contributed to conceptualization of the study. S.C. and T.Z. performed additional biochemical experiments. S.W. and J.M. designed and performed simulation experiments and analyzed the related data. R.P. and S.W. helped with creation and validation of plasmids. D.J., and J.A. and MD.R generated the plasmids for FLAG-FUS expression and U2OS GFP-FUS cell line generation. D.J. and J.A. performed the cell-based assays in cell lines. M.B. and B.S.S. performed the microrheology. A.B. performed the fly experiments, including crosses, western blotting, and data quantification in *Drosophila* with supervision, analysis, and guidance from U.B.P. S.K. designed and planned fly experiments, including data quantification and filter trap assays, compiled results, and wrote the Methods for *Drosophila*. E.N.A. carried out dissections and confocal imaging in *Drosophila*.

## Conflict of Interest Statement

The authors declare no conflict of interest.

## STAR Methods

### CONTACT FOR REAGENT AND RESOURCE SHARING

Further information and requests for materials may be directed to and will be fulfilled by the Lead Contact.

### EXPERIMENTAL METHOD AND SUBJECTS DETAILS

#### Plasmids

For eukaryotic expression, pcDNA6F-hsFUS (Plasmid ID: C19) was generated previously^93^ and pcDNA6F-12P-hsFUS was generated by digesting pcDNA6F-hsFUS with XhoI (NEB, #R0146) and BmgBI (NEB, #R0739) and replacing the excised XhoI-BmgBI fragment with a gene-synthesized (GeneArt, Life Technologies) fragment encoding the human FUS N-terminal region containing the 12 additional prolines. For genome editing, pMK-HDR-eGFP-mmFUS-WT-BSD (Plasmid ID: I123) is described in^37^. To create pMK-HDR-eGFP-mmFUS-12P-BSD (Plasmid ID: I132), pMK-HDR-eGFP-mmFUS-WT-BSD was digested with BsrG1I (NEB, #R3575) and the excised fragment replaced with a gene-synthesized fragment (GeneArt, Life Technologies) encoding the mouse FUS N-terminal region containing the 12 additional prolines. To create pMK-HDR-eGFP-mmFUS-G507Vfs-BSD (Plasmid ID: I126), the β-globin chimeric intron region was amplified (Primer IDs: DJ852 and DJ853) from pMK-HDR-FUSKI-G515Vfs-BSD (Plasmid ID: I115) described in^37^. The eGFP-mmFUS coding region was amplified (Primer IDs: DJ854 and DJ855) from pcDNA6-eGFP-GSG15-mmFUS-G507Vfs (Plasmid ID: C82). The two fragments were then fused by PCR (Primer IDs: DJ852 and DJ855) and cloned into the BamHI (NEB, #R3136) and NotI (NEB, #R3189) sites of pMK-HDR-FUSKI-G515Vfs-BSD (Plasmid ID: I115). Finally, pMK-HDR-eGFP-mmFUS-P12-G507Vfs-BSD was generated by amplifying the mutated NLS from pMK-HDR-eGFP-mmFUS-G507Vfs-BSD (Plasmid ID: I126) by PCR (Primer IDs: JM96 and mdr188), and cloning into NdeI (NEB, #R0111) and NotI (NEB, #R3189) digested pMK-HDR-eGFP-mmFUS-12P-BSD.

#### Oligonucleotides

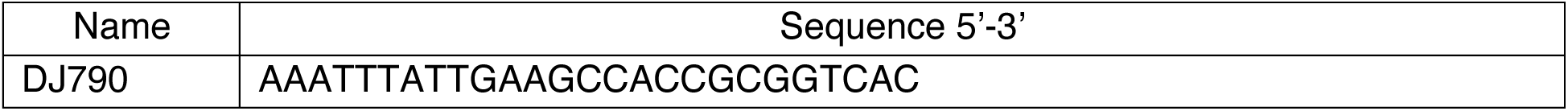
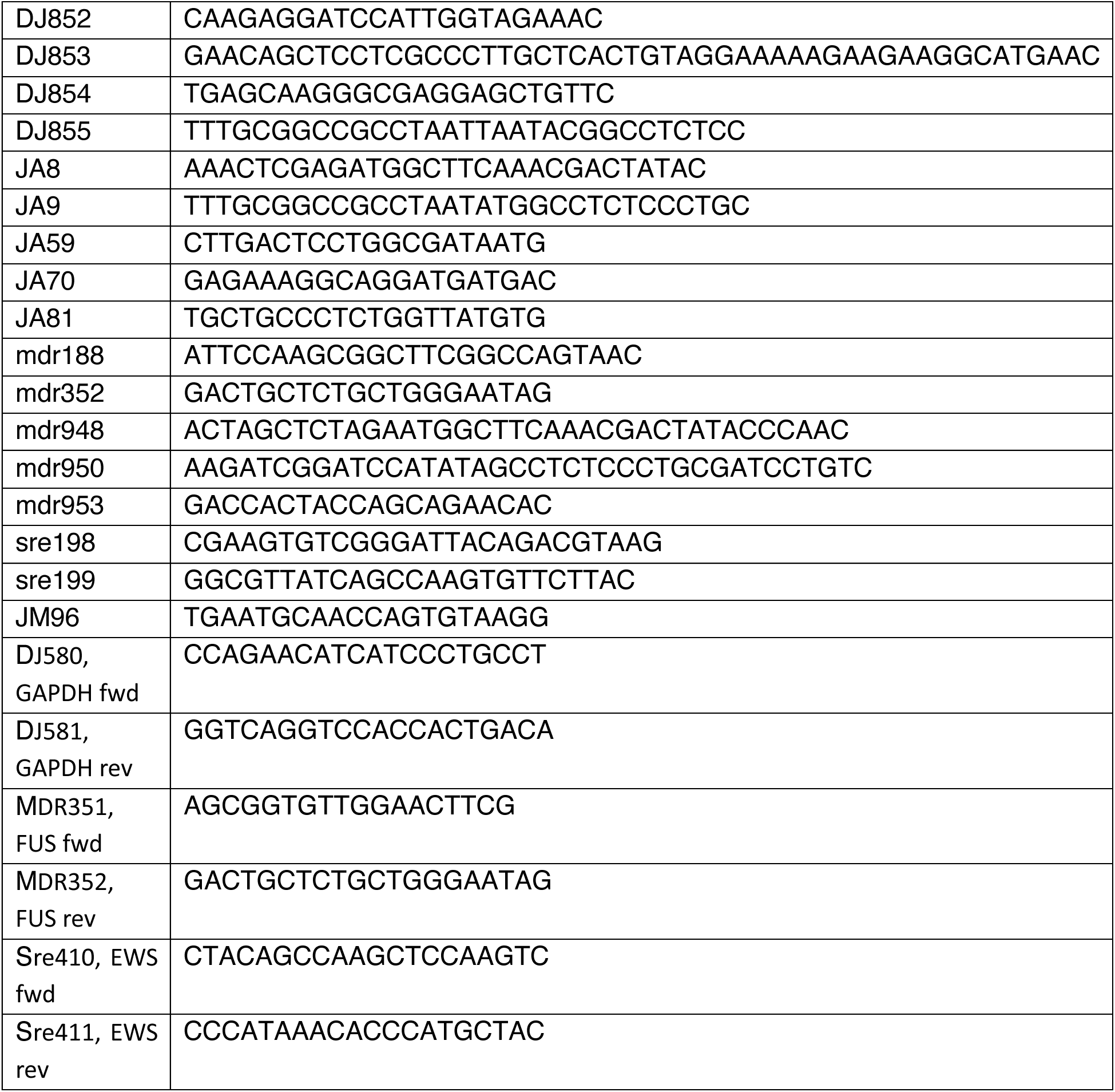

#### Cell Lines

U2OS and HeLa cells were purchased from ECACC (Acc N° 92022711 and Acc N° 93021013) and maintained at 37°C and 5% CO2 in DMEM/F12 GlutaMAX (Gibco, #10565018) supplemented with 10% (v/v) fetal bovine serum (FBS) (PAN-Biotech, #P30-3031) and 100 U/ml (1% v/v) Penicillin-Streptomycin (Gibco, #15140122). Cells were passaged using TrypLE Express (Gibco, #12605010) unless stated differently.

#### Drosophila lines

UAS FUS-WT, FUS-P525L, FUSWT-12P and FUS-P525L-12P lines were generated by Best Gene Inc by site-specific insertion of the human transgene in BDSC#8622 attp2 injected stock using the pUAST-attP2 vector. The fly stocks were maintained on a standard dextrose medium in a 12-hour light/dark cycle incubator. All the fly images were taken using a Leica M205C dissection microscope equipped with a Leica DFC450 camera.

## Methods

### Bacterial protein expression and purification

BL21 Star DE3 cells expressing 6xHis-tagged FUS LC, LC-RGG1, and MBP-tagged FUS full-length were grown in M9 minimal media (27 mM NaCl, 22 mM KH_2_PO_4_, 51 mM Na_2_HPO_4_×7H_2_O, 1 mM MgSO_4_, 1% v/v MEM vitamin solution 100X, 0.2%v/v solution Q (composed of 8%v/v 5M HCl, 0.5% w/v FeCl_2_×4H_2_O, 0.018%w/v CaCl_2_×2H_2_O, 0.0064% w/v H_3_BO_3_, 0.0018% w/v CoCl_2_×6H_2_O, 0.0004% w/v CuCl_2_×2H_2_O, 0.034% w/v ZnCl_2_ and 0.004% w/v Na_2_MoO_4_×2H_2_O), 4 g of glucose, and 1 g ammonium chloride) using ^15^N labelled ammonium chloride as the sole nitrogen source. Cells were grown at 37 °C with 200 rpm shaking until they reached an OD at 600 nm of 0.8. Protein expression was induced by addition of 0.5 mM of IPTG (isopropyl β-D-1-thiogalactopyranoside) and allowed to express for 4 hours at 37 C and 200 rpm shaking. For the LC and LC-RGG1 variants: Cells were resuspended in 20 mM sodium phosphate pH 7.4, 300 mM sodium chloride. Lysis was performed using a Fisherbrand sonicator using a square wave sonication profile for 30 second intervals and 20 minutes total sonication time and 75% of the maximal amplitude. Cell lysate was then centrifuged at 47,855 *g* for 30 minutes. The cell lysate supernatant was discarded and the inclusion body was solubilized in 20 mM sodium phosphate, pH 7.4, 300 mM sodium chloride buffer containing 8 M urea and 10 mM imidazole. This solution was then centrifuged at 47,855 *g* for 30 minutes and the supernatant was collected and filtered using a 0.45 micron syringe-driven filter unit.

For the MBP-tagged full length variants: Cells were resuspended in 20 mM sodium phosphate pH 7.4, 10 mM imidazole, and 1 M sodium chloride. Lysis was performed using similar methods as the LC and LC-RGG1 sequences. Once lysed, the cell lysate was centrifuged at 47,855 *g* for 30 minutes and the supernatant was collected and filtered using a 0.22 micron syringe-driven filter unit.

LC and LC-RGG1 purification: The resolubilized inclusion body was loaded onto a 5 mL immobilized nickel-affinity column (IMAC; Cytiva) attached to a Bio-Rad NGC. The protein was then eluted using a gradient of 10 – 300 mM imidazole. Fractions containing the protein were concentrated to 1 mL using an Amicon 3 kDa MWCO centrifugal filter unit. The concentrated protein was then diluted 10× into sodium phosphate pH 7.4 buffer containing TEV protease and TEV cleavage was allowed to proceed over night at room temperature on a gently rocking table. Once cleaved, urea, imidazole, and sodium chloride were added to the solution to bring final solution components concentrations to 20 mM sodium phosphate, 300 mM sodium chloride, 10 mM imidazole, and 8 M urea. This solution was loaded onto the 5 mL nickel IMAC column (HisTrap HP, Cytiva) to separate the cleaved 6xHis tag, TEV protease, and the protein of interest. Fractions containing the protein of interest were buffer exchanged into 20 mM CAPS pH 11.0 until the residual urea concentrations were < 10 μM urea in the final concentrated stock solutions. These samples were then flash frozen and stored at -80 °C for later use.

For the full-length variant: The cell lysate containing MBP-FUS full-length were loaded onto a 5 mL nickel IMAC column and then eluted using a gradient of 10 – 300 mM imidazole. Fractions containing the protein were pooled and dialyzed over night at against 20 mM sodium phosphate pH 7.4, 1 M sodium chloride buffer. The sample was then spin concentrated using an Amicon 10 kDa MWCO centrifugal spin filter unit to 10 mL and loaded onto a Superdex 200 26/600 column equilibrated in 20 mM sodium phosphate pH 7.4 and 1 M sodium chloride. Fractions corresponding to MBP-FUS full-length were collected, pooled, and concentrated to ∼ 400 μM and flash frozen for later use.

### NMR

#### Relaxation Analyses

Dilute phase NMR studies were performed on 75 μM samples of FUS LC wild-type and 12P prepared by 10-fold dilution from the 20 mM CAPS pH 11.0 storage buffer to final sample conditions of 20 mM MES pH 5.5, 150 mM sodium chloride, 10%v/v storage buffer, and 4.2% deuterium oxide lock solvent prepared in a 5 mm NMR tube. ^1^H,^15^N HSQC spectra were collected with 3072 real and imaginary points in the direct dimension and 256 indirect dimension points. Spectral widths for the ^1^H and ^15^N channels were set to 10.0 and 20 ppm, respectively. 8 scans were collected for each indirect dimension point. CCPNMR was used to visualize the HSQC spectra.

Heteronuclear ^1^H-^15^N relaxation parameters were acquired using standard NMR pulse sequences (hsqct1etf3gpsitc3d, hsqct2etf3gpsitc3d, hsqcnoef3gpsi). *R*_1_ and *R_2_* for the dilute phase samples were collected with 4096 direct and 400 indirect points. These experiments utilize spectral widths of 10.5 and 20 ppm for the ^1^H and ^15^N channels, respectively and carrier frequencies of 4.7 and 117 ppm, respectively. These relaxation studies were performed using interleaved relaxation times to minimize effects of sample heating. For the *R*_2_ experiments, relaxation delay values of 16.9, 270.4, 185.9, 33.8, 118.3, 84.5, 169 ms were sampled using a 556 Hz CPMG field. For the *R*_1_ measurements, delay values of 100, 1000, 200, 800, 300, 600, 400 ms were used. For the ^15^N heteronuclear NOEs, interleaved steady-state NOE and no-NOE control spectra were collected with 4096 indirect points and 354 total indirect points. For the NOE experiments a 5 s recycle delay and a 100 ms mixing time were used. Carrier frequencies of 4.7 and 117 ppm were used for ^1^H and ^15^N ppm, respectively. Relaxation values were determined using NMR-PIPE to fit the relaxation single exponential decay curves 𝐼 = 𝐼_0_ × 𝑒^(-*R*^*^it^*^)^, where *I* is the signal intensity, *I*_0_ is the signal intensity at short relaxation delay time, *R_i_* is the transverse or longitudinal decay rate constant, and *t* is the experimental relaxation delay time. Relaxation profiles were plotted using GraphPad Prism v10.

Samples to measure NMR relaxation values in the condensed phase were created by sedimenting condensed phase LC-RGG1 and carefully transferring the material into a 3 mm tube. Phase separation was induced by diluting FUS LC-RGG1 10-fold from its storage buffer (20 mM CAPS pH 11.0) into 50 mM MES pH 5.5, 150 mM sodium chloride and 10% v/v ^2^H_2_O for the NMR lock. Standard NMR heteronuclear relaxation pulse sequences were used (hsqct1etf3gpsitc3d, hsqct2etf3gpsitc3d, hsqcnoef3gpsi). Spectra were acquired with carrier frequencies of 4.7 and 117 ppm and spectral widths of 10.5 and 30 ppm for the direct and indirect dimensions, respectively. The total number of points for the direct and indirect dimensions were 4096 and 256 points, respectively. ^15^N transverse relaxation values, *R*_2_, were measured using a 7-point interleaved relaxation delay values of 16.9, 270.4, 185.9, 33.8, 118.3, 84.5, 169 ms and a 556 Hz CPMG field while ^15^N longitudinal relaxation values, *R*_1_, were measured using a 7-point interleaved relaxation delay values of 100, 1000, 200, 800, 300, 600, 400 ms. A 5 s recycle delay and 100 ms mixing time was used for the ^15^N NOE similarly used carrier frequencies of 4.7 and 117 ppm for direct and indirect dimensions, respectively, and spectral widths of 10.5 and 30 ppm, respectively. Total number of points for the direct and indirect dimensions were 4096 and 256, respectively. NOE experiments were collected as interleaved steady-state NOE and no-NOE control experiments.

#### Diffusion

NMR diffusion coefficients were measured on dilute FUS LC and condensed phase LC-RGG1 samples using standard NMR experiments (ledbpgppr2s) using the sample samples as described above. For the dilute phase, 16 linear gradient strengths ranging from 2 – 98% of encode/decode gradient were optimized for a fixed gradient length of 1.5 ms. The total diffusion time (Δ) was 150 ms. For the condensed phase, 16 linear gradient strengths ranging from 2 – 98 % encode/decode gradient were optimized for a fixed gradient length of 7 ms. The total diffusion time (Δ) was set to 2 s. Data was processed using Topspin 4.3.0 by integrating peaks corresponding to the FUS LC protein and data was fit to the equation, 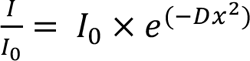, where, *I/I_0_*, is the measurement of the fraction of coherent NMR signal remaining after diffusion during application of a gradient, A, is a constant, *D* is proportional to the diffusion constant, and, *x*, is the gradient strength percentage.

### Circular Dichroism

Solutions of FUS LC wild-type and 12P were prepared to be 5 μM FUS LC in 20 mM sodium phosphate pH 7.4 with 150 mM sodium chloride. CD spectra were acquired at 25 °C over the wavelengths 195 – 280 nm using a Jasco J-815 CD spectrometer. Spectra were acquired as the mean of technical triplicate runs.

### Phase Separation Quantification

#### Salt dependent phase diagram

The salt dependent phase diagrams of FUS LC or LC-RGG1 was measured as previously reported^14^. In brief, FUS LC or LC-RGG1 was diluted 10× from its storage buffer (20 mM CAPS, pH 11.0) to a final concentration of 300 or 60 μM, respectively, and a final volume of 30 μL using 20 mM HEPES pH 7.0 (pH adjusted using 1M BIS-TRIS) and containing varying molar quantities of sodium chloride. Samples were mixed well and then centrifuged for 20 minutes at 17,000 *g*. The absorbance at 280 nm was measured for the supernatant of each sample. These experiments were performed with n=3 technical replicates and at least N=2 biological replicates.

#### Cloud Point Measurements

Measurements of the temperature dependent phase diagram of FUS LC-RGG1 wild-type and 12P variants were performed by diluting the LC-RGG1 constructs 10× from their storage buffer (20 mM CAPS pH 11.0) into 20 mM HEPES pH 7.0 buffer containing 150 mM sodium chloride that was equilibrated on a hot plate set to 90 degrees Celsius and immediately transferred to a 96 well plate similarly equilibrated to 90 degrees Celsius using a hot plate. Final protein concentrations of 60, 80, and 120 μM LC-RGG1 were used for analysis and samples were visually confirmed to be absent of any phase separation. The 96-well plate was then loaded into a Cytation 5 plate reader that has been equilibrated at 65 degrees Celsius and the plate was allowed to equilibrate to 65C. Turbidity measurements at 350 nm were collected for sensitive detection of phase separation. The temperature was lowered in 1-degree Celsius increments and allowed to equilibrate for 1 minute between each temperature step. The temperature was lowered to a minimum value of 25 degrees Celsius. The resulting data was processed using an in-house Matlab script (Matlab 2024b) that fits the inflection point of the turbidity at 350 nm from its initial value to its maximal value by fitting the curves to the equation, 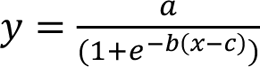, where, *y,* is the measured absorbance at 350 nm, *a,* is the maximal absorbance value, *b,* is the minimal absorbance value, and, *c,* is the temperature in which the half maximal absorbance is observed. For each biological replicate, n=8 technical replicates were individually fit to determine the temperature in which the turbidity reached its half-maximal value, and the average of the fitted parameter was interpreted as the cloud point value for that biological replicate. Then, the mean and standard deviation of N=3 biological replicates was calculated.

#### Direct measurement of Condensed Phase concentration

Stock samples (stored in 20 mM CAPS pH 11.0) were concentrated to ∼5 mM and diluted 10× into 20 mM HEPES pH 11.0 with 150 mM sodium chloride. The samples were then centrifuged at 17,000 *g* for 10 minutes. Triplicate measurements were then performed by pipetting 10 μL of condensed phase with a positive displacement pipette and diluting it 1000-fold into 20 mM HEPES pH 7.0, 150 mM sodium chloride buffer containing 8 M urea.

#### Turbidity

Concentrated MBP FUS FL WT and 12P (stored in 20 mM sodium phosphate, 1 M NaCl, pH 7.4) was diluted to 100 μL to yield 20 µM FUS in a final solution condition of 20 mM HEPES 150 mM NaCl pH 7.0. 2 μL of concentrated TEV protease was added to each sample to cleave the solubilizing MBP tag and induce phase separation. Samples were equilibrated for 30 minutes on the benchtop to allow cleavage to occur. n=8 technical replicates were performed by transferring individually prepared samples to a Greiner 96 well half-area microplate. Turbidity was evaluated by measuring the absorbance at 600 nm in a Cytation 5 Cell Imaging Multi-Mode Plate Reader. Student’s t-test was performed to test for differences between the wild-type and 12P samples using GraphPad Prism 10.

### Microscopy

#### Fluorescence microscopy

Phase separation and droplet morphology was assessed by diluting FUS LC or LC-RGG1 ten-fold to a final concentration of 300 μM or 60 μM, respectively using a buffer composed of 20 mM HEPES pH 7.0 (adjusted with 1 M Bis-Tris), 150 mM sodium chloride, and 22 μM fluorescent dye ThioflavinT. The samples were mixed thoroughly before 20 μL was transferred to a glass coverslip for imaging using a Nikon Ti2 DIC microscope using a 20× objective with 1.5× digital enhancement.

For the full-length FUS (stored as MBP-FUS), samples were prepared by 6.7× dilution from the storage buffer (20 mM NaPi, pH 7.4, 1M sodium chloride) to a final protein concentration of 60 μM using 20 mM HEPES buffer, pH 7.0 (adjusted using 1 M Bis-Tris), and 22 μM Thioflavin T. 2 μL of TEV protease was added to each sample and samples were then mixed well and incubated for 30 minutes at room temperature before 20 μL of each sample was transferred to a glass coverslip and imaged using a 20× objective with 1.5× digital enhancement.

#### DIC Microscopy

For samples where phase separation in response to treatment with PEG 8000 was examined, FUS LC-RGG1 was diluted 10× from its storage buffer (20 mM CAPS, pH 11.0) to a final protein concentration of 15 μM into 20 mM HEPES, pH 7.0 (adjusted with 1 M Bis-Tris), 150 mM sodium chloride, and 5%w/v PEG. Samples were mixed thoroughly before 20 μL was transferred to a glass coverslip and imaged using a Nikon Ti2 DIC microscope using a 20× objective with 1.5× digital enhancement.

For samples where aggregation was examined over time, independent samples for each time point were generated by diluting FUS LC 10× from its storage buffer using 20 mM HEPES pH7.0 (adjusted with BIS-TRIS), 150 mM sodium chloride to a final concentration of 300 μM and a total volume of 100 μL. All samples were then placed on an Eppendorf Thermomixer C equilibrated at 25 C and allowed to shake at 1200 rpm for the desired duration. Samples corresponding to each time point were then removed from the thermomixer and mixed thoroughly before 20 μL was transferred to a glass slide and visualized using a Nikon Ti2 DIC microscope with a 20× objective using 1.5× digital enhancement.

For monitoring the aggregation of the full-length sequences, concentrated stock solutions of FUS were diluted 6.7× into 20 mM HEPES, pH 7.0 (adjusted with 1 M Bis-Tris) to a final protein concentration of 60 μM and a total volume of 100 μL. Independent samples were used for each desired time point. Then, 2 μL of TEV protease was added and the samples were thoroughly mixed and allowed to equilibrate at room temperature for 30 minutes before loading them onto a Eppendorf thermomixer equilibrated at 25 C and shaken at 1200 rpm for the specified duration. At the respective time point, each sample was removed from the mixer and mixed thoroughly before 20 μL was transferred to a glass cover slip for imaging.

### Simulations

#### All-Atom Simulations Protocol and Trajectory analysis

Wild-type simulation trajectories were obtained from previous work^14^. Single-chain simulations of the 12P variants were performed using the same protocol—employing the AMBER03ws force field with the TIP4P/2005s water model and updated NaCl parameters—to ensure direct comparability of results^94,95^. All the production runs were performed under a canonical ensemble (NVT) at 300 K.

Protein–protein pairwise contacts and average intrachain distances (Rij) were calculated using custom scripts built on MDAnalysis 2.5.0^96,97^. Contact was defined as present when any heavy atoms from two residues were within 4.5 Å of each other. Pairwise contacts were quantified by summing all heavy atom interactions between residue pairs, excluding those involving residues within a five-residue sequence proximity. R_ij_ values were computed based on the positions of Cα atoms.

### Thioflavin T fluorescence / Aggregation assay

For the LC samples: n=3 independent samples for each time point comparing FUS LC wild-type, S→G, or 12P, or comparing FUS LC wild-type, 4P, 12P, and 16P, were prepared by 10× dilution from 20 mM CAPS pH 11.0 storage buffer into phase separating conditions of 300 μM FUS LC in 20 mM HEPES pH 7.0, 150 mM sodium chloride, and 20 μM Thioflavin T, and then mixed well. These samples were then affixed to an Eppendorf Thermomixer and subjected to orbital shaking at 1200 rpm for the desired total duration. At the end of each time point, the samples were removed from the thermomixer and mixed thoroughly before transferring to a 96-well plate that was loaded into a Cytation 5 plate reader. The fluorescence at 480 nm was measured. Then, a representative sample was transferred from the plate to a coverslip for imaging by DIC microscopy.

### Microrheology

#### Video particle-tracking microrheology

Yellow–green carboxylate-modified polystyrene beads (500-nm diameter; FluoSpheres, Invitrogen) were used for video particle-tracking (VPT) microrheology measurements. Both WT and 12P samples were prepared at the final concentration of ∼60 µM for microrheology by diluting the stock solution. Before imaging, the samples were kept at 75^0^C for 10 mins so that there was no droplet assembly before the start of the experiment. Next, 120 μl protein sample was removed, mixed with polystyrene beads and transferred to the 384-well plate (#1.5 high-performance cover glass, Cellvis) to initiate phase separation at room temperature. The samples were then incubated in a well plate at room temperature for 5 minutes followed by the centrifugation step at 400x*g* for 3 mins to form a condensate layer or larger-sized droplets (>20 µm in diameter); the purpose of this step was to avoid boundary effects and prevent flow of the condensates.

Next, epifluorescence video imaging was performed on a Zeiss Axio Observer 7 inverted microscope equipped with an Axiocam 702 monochrome sCMOS camera (Zeiss), employing a x63/1.4-NA plan-apochromatic oil-immersion objective, with fluorescence excitation using a 475-nm light-emitting diode (Colibri 7; Zeiss). The microscope focus was adjusted to the midsection of the protein sample for VPT acquisition. Data collection was initiated at 1 hr, 24hrs, 48hrs and 72 hrs timepoint, with incubation of sample at 20^0^C in between measurements. Imaging was conducted at room temperature (19–20 °C). Videos of the tracer beads diffusing within the condensate were collected at 200 frames per second for 2,000 frames and saved for further analysis. For each variant, two independent samples were made on different days, and four to five videos were collected from each sample. MSD and slope data presented in Fig.3D are the averages of these two independent trials (*n* = 8-9 for the data shown).

The TrackPy particle tracking code was used to analyze the collected videos, starting with extracting particle trajectories. The MSD was calculated from the trajectories of individual beads, followed by calculating the ensemble-average MSD. To remove the static error from the MSD curves, we corrected the ensemble-average MSD by subtracting the noise floor from the MSD curves. In general, the ensemble-average MSD often scales as a power law with lag time *τ*, as given by MSD (*τ*) = 2dD*τ*^α^ here *d* is the number of dimensions (here *d* = 2, because data collection and analysis were conducted in the *x–y* plane), *D* is the diffusion coefficient, and *α* is the diffusivity exponent. The *α* values were extracted by fitting the average MSDs from 0.1 to 1 s to the model outlined above. For a purely viscous fluid, the diffusivity exponent *α* approaches 1 at long lag times whereas *α* < 1 signifies a sub-diffusive behavior or deviation from purely viscous behavior.

### Fluorescent Protein Partitioning

#### Alexa488-FUS / Alexa488-RNAPII CTD

Fluorescent-tagged FUS LC and RNA Pol II CTD were generated by NHS ester labelling and performed by separately diluting the proteins to 100 μM in 20 mM sodium phosphate buffer pH 7.4 containing 150 mM sodium chloride and a 5× molar excess of Alexa488 (DyLight 488 NHS-Ester). This solution was incubated at room temperature overnight. Free dye was removed by buffer exchanging into fresh phosphate buffer using a 3 kDa molecular weight cut off centrifugation filter unit (Amicon) until the flow-through contained non-detectable amounts of fluorescent label.

Partitioning experiments were performed by adding 2 μM of the fluorescently tagged protein to solutions of 300 μM FUS LC wild-type or the 12P in 20 mM HEPES pH 7.0 buffer containing 150 mM sodium chloride. The FUS LC was diluted ten-fold from its CAPS pH 11.0 storage buffer to ensure that the total concentration of CAPS in the final sample was 10% v/v. Samples were mixed thoroughly and the condensed phase was sedimented by centrifugation at 17,000 *g* for 20 minutes. The fluorescence of the supernatant was then measured. To determine the fraction of alexa488 labelled protein that partitioned into the FUS LC condensed phase, these samples were compared to a control sample containing 2 μM fluorescently labeled protein and no FUS LC condensed phase under identical buffer conditions (10% CAPS pH 11.0, 90% 20 mM HEPES pH 7.0 150 mM sodium chloride) and the fraction was obtained by dividing the test sample by the control sample. The uncertainty was propagated using the equation, 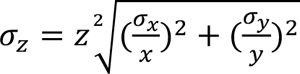, where *x* corresponds to the mean values of the raw fluorescence units of the test sample (n=3), *y* corresponds to the mean of the raw fluorescence units of the control sample, *z* corresponds to the ratio of *x* and *y*, *σ* corresponds to the standard deviation of the respective subscript.

### FUS auto- and cross-regulation assay

The auto and cross-regulation assay was essentially performed as described in^1^. In brief, HeLa cells were grown to 80% confluency in six-well plates and transfected with 1000 ng pcDNA6F-FUS-WT or pcDNA6F-FUS-12P using Transit-LT1 (Mirus, #2304) according to the manufacturer’s recommendations. Non-transfected cells (NTC) served as control. Cells were split into two six-well plates after 24 hours and harvested 48 hours post-transfection: One well in 600 µl RNA lysis buffer (Agilent, #400800), one well in 100 μl RIPA buffer (Thermo, #89900) containing 1× Halt™ Protease Inhibitor Cocktail (Thermo, #87786). Western blot analysis was performed as outlined below, with the following antibodies: Mouse anti-FLAG M2 (Sigma, #F3165), Rabbit anti-FUS serum 6 (home-made, as previously described^93^), Mouse anti-GAPDH (Santa Cruz, #sc32233). RNA was isolated with the Agilent Absolutely RNA miniprep kit (Agilent, #400800) according to the manufacturer’s protocol and RNA concentrations were measured using a NanoDrop™ One Microvolume UV–Vis Spectrophotometer (Thermo Fisher, #ND-ONE-W). Subsequently, cDNA was generated from 1 µg of RNA using LunaScript RT SuperMix (NEB, #E3010) according to the manufacturer’s instructions. Following dilution to a concentration of 8 ng/µl, qPCR was performed with PowerUp SYBR mix (Applied Biosystems, #A25742) on a Rotor-Gene 6000 Q thermal cycler (Qiagen, #9001550). All samples were measured in technical triplicates. For each reaction, 4 μl of cDNA were amplified in a total volume of 20 μl containing 600 nM of both forward and reverse primers (Primer IDs: GAPDH: DJ580, DJ581, FUS: mdr351, mdr352, EWS: sre410, sre411). Note that the FUS primers are specific to the endogenous transcript and do not bind to exogenous FLAG-FUS cDNA. Gene expression analysis was performed using the ΔΔCt method, and statistical significance was assessed using an unequal variances *t*-test applied to the ΔΔCt values.

### Genome editing

Genome editing of U2OS lines was performed as described previously^37^. In brief, cells were transfected with a Cas9 and gRNA expressing plasmid targeting *FUS* intron 1 (TGGATGTCCACCAAGACCT), alongside homology direct repair (HDR) gene replacement matrices encoding the different GFP-mmFUS versions using *Trans*IT-LT1 Transfection Reagent (Mirus Bio, #MIR 2300) according to manufacturer’s instructions. 48h post-transfection, cells were selected with 10 μg/ml blasticidin S HCl (Gibco, #A1113903) for 5 days. After selection, cells were dissociated and transferred as single cell suspension to 15 cm 0 cell culture dishes. Colonies with cells showing homogeneous GFP signal were isolated using cloning cylinders and transferred to single wells of cell culture plates for further validation by WB using α-FUS (#sc47711) and α-GFP (**#**A-11122) antibodies and by PCR as described previously^37^.

### DNA Damage Response

All DNA damage response assays were conducted, and data were recorded and analyzed as detailed^37^. In short, for microirradiation assays, cells were seeded in poly-d-lysine (Gibco, #A3890401) coated glass-bottom culture dishes, pre-sensitized with bisBenzimide H 33342 trihydrochloride (Hoechst, Sigma, #B2261), and media changed to DMEM Fluorobrite (Gibco, #A1896701) supplemented with 2% FBS. Live cells were microirradiated at 37°C and in a 5% CO_2_ atmosphere using a 40× water immersion lens and the 405 nm laser in a Nikon AX inverted confocal microscope. For chemical-induced DNA damage assays, cells were seeded in PhenoPlate cell imaging microplates, incubated overnight at 37°C and treated for 1h with 5 µM etoposide (ETO, #CAY12092) or 50 nM calicheamicin γ1 (CAL, #CAY40830) before fixing and high-throughput imaging using an Opera Phenix Plus high-content screening system (Revvity). Non-treated controls contained 0.025% (v/v) DMSO (Apollo Scientific, #67-68-5). Calicheamicin-induced FUS phosphorylation western blots were done using extracts from cells seeded in 6-well cell culture dishes and treated with increasing concentrations of calicheamicin γ1 for 1h, or with 50 nM for different durations, as indicated in the figure.

### eGFP-FUS paraspeckle localization

Imaging of paraspeckles in U2OS cells and analysis of eGFP-FUS nuclear localization and foci number were carried out as specified previously^37^. Briefly, after high-throughput imaging of fixed cells stained with PSCP1 antibodies, eGFP-FUS foci number was counted using automated analysis pipelines in Harmony software (Revvity, v4.9.2137.273), and co-localization with PSCP1 was analyzed by comparing the fluorescence intensity profiles along the white line indicated in the images using FIJI (v.2.0.0), normalising the intensities to the maximum value for each channel scaled from 0 to 1.

### eGFP-FUS cytoplasmic assemblies formation

Cells were seeded in PhenoPlate cell imaging microplates, incubated overnight at 37°C. Then, where indicated, cells were incubated at 44°C (heat-shock, HS) for 2h and recovered at 37°C in the presence of 50 μM NaASO_2_ (Sigma, #106227). Alternatively, cells were transfected with high (Sigma-Aldrich, Cat#P9582) or low molecular weight Poly(I:C) (InvivoGen, #tlrl-picw) using Lipofectamine 2000 (Invitrogen) and incubated for 4h. After Poly(I:C) treatment, cells were washed with media and recovered in fresh media at 37°C for 2, 4 and 8h. After the different treatments or recovery timepoints, cells were fixed with 8% PFA (1:1, v/v) (Thermo Scientific, #047347.9M) and processed for ICC and further high-throughput analysis.

### Immunocytochemistry (ICC)

ICC of cell-based assays was generally done as described in^37^. Shortly, cells were washed with PBS and fixed with PFA 4% (Thermo Fisher, #J19943.K2) at room temperature (RT) for 15 min. Cells were washed and permeabilized with 1x TBS containing 0.5% (v/v) Triton-X (PanReac, #A4975,0500) and 6% (w/v) bovine serum albumin (BSA, Sigma, #A9418) for 30 min. Next, cells were incubated overnight at 4°C with primary antibodies diluted in 1x TBS with 0.1% (v/v) Triton-X 6% and (w/v) BSA. Next day, cells were washed three times and incubated with Alexa Fluor-conjugated secondary antibodies and DAPI (Life Technologies, #834650) for 1h at RT, washed three times and kept at 4°C until imaging.

### High-throughput screening image acquisition and analysis

Imaging of all GFP-FUS foci, Cajal bodies/SMN foci, paraspeckles, and DDR was done using an Opera Phenix Plus high-content screening system (Perkin Elmer) using 40× or 60× water-immersion lenses in confocal mode. Z-stacks were taken every 0.5 μm covering the whole 3D area of cells. Acquisition for each channel was kept separated throughout all scans to avoid crosstalk between different fluorescent signals. Quantitative analysis of the different readouts was done in the maximum projection of all Z-stacks

### Developmental assays

Eclosion assays to assess developmental defects in flies were conducted as previously described^81^. UAS-FUS and 12P flies were crossed with the motor neuron–specific driver D42-Gal4 at 28 °C, and eclosion of adult flies from puparia was monitored over a 4-day period (n=3). The total numbers of eclosed adults and puparia (both eclosed and non-eclosed) were quantified, and the eclosion rate (adult/pupa ratio) was calculated for each condition and normalized to the corresponding controls.

Similarly, flies exhibiting the crumpled wing phenotype were quantified for each cross, and the wing defect frequency was calculated as the ratio of crumpled to straight wings.

### Motor-function assays

RING assay: UAS-FUS and 12P flies were crossed with the motor neuron–specific driver D42-Gal4, female progeny was collected at day 1. Locomotor activity was assessed on day 7 using the rapid iterative negative geotaxis (RING) assay as described previously^83,98^. The flies were tapped to the bottom of vials, and climbing was recorded for 30 s (n=10). Locomotor performance was quantified either as climbing velocity (distance per unit time) or as the percentage of flies reaching 10 and 5cm mark within 10 and 5 s, respectively. Data was normalized to w1118 control flies.

### Life span

F1 flies expressing FUS-WT, FUS-P525L, FUS-WT-12P, or FUS-P525L-12P (20 flies per vial; n = 50) were monitored daily, and mortality was recorded. Survival was analyzed using Kaplan–Meier plots. Control strain (w1118) flies crossed with D42 were used as controls.

### Eye degeneration

UAS-FUS and 12P flies were crossed with the eye-specific driver GMR-Gal4 at 29 °C. F1 female progeny were collected on day 4, and eye morphology was imaged using a Leica M205C microscope. External degeneration was quantified using a published scoring system^80,83,99^, with scores assigned according to the extent of abnormal bristle orientation, retinal collapse, ommatidial fusion, pitting, and disorganization of the ommatidial array.

Statistical comparisons between genotypes were performed using Student’s t-test.

### TUNEL assay

Third-instar larval brains from F1 flies expressing FUS-WT, FUS-P525L, FUS-WT-12P, or FUS-P525L-12P were fixed in 4% paraformaldehyde. Neuronal apoptosis was detected using the In Situ Cell Death Detection Kit, Fluorescein (Sigma), according to the manufacturer’s instructions. Brains were mounted in DAPI-Fluoroshield (Sigma, F6182), and images were acquired using a Nikon A1 Eclipse Ti confocal microscope.

### Western blotting

Nine fly heads were collected, snap-frozen, and homogenized in RIPA buffer (150 mM NaCl, 1% NP-40, 0.1% SDS, 1% sodium deoxycholate, 50 mM NaF, 2 mM EDTA, 2 mM DTT, 0.2 mM Na₃VO₄, 1× protease inhibitor). Lysates were sonicated, centrifuged, and supernatants boiled in Laemmli buffer. Separated proteins in 4–12% NuPAGE Bis-Tris gels were transferred to nitrocellulose membranes (iBlot2, Invitrogen) and probed overnight with rabbit anti-FUS and mouse anti-tubulin, followed by IRDye 680D anti-mouse and DYLight 800 anti-rabbit secondary antibodies. Blots were imaged on an Odyssey CLx system. Experiments were performed in triplicate (n = 9), and protein levels were quantified using Student’s t-test.

For filter retardation assay, 20 μg of total protein were filtered on a 0.2-μm cellulose acetate membrane (Whatman GE Healthcare, GEH10404180) using a Bio-Dot SF Microfiltration Apparatus (Bio-Rad Laboratories, 1703938) (n=6).

### Data and Software Availability

All data are available upon reasonable request. Bacterial expression plasmids will be deposited to Addgene.

## Notes

### Competing Interest Statement

The authors have declared no competing interest.

