## Supplementary Figures for "Breaking β-sheets in FUS prion-like domain preserves phase separation and function but prevents aggregation and toxicity"

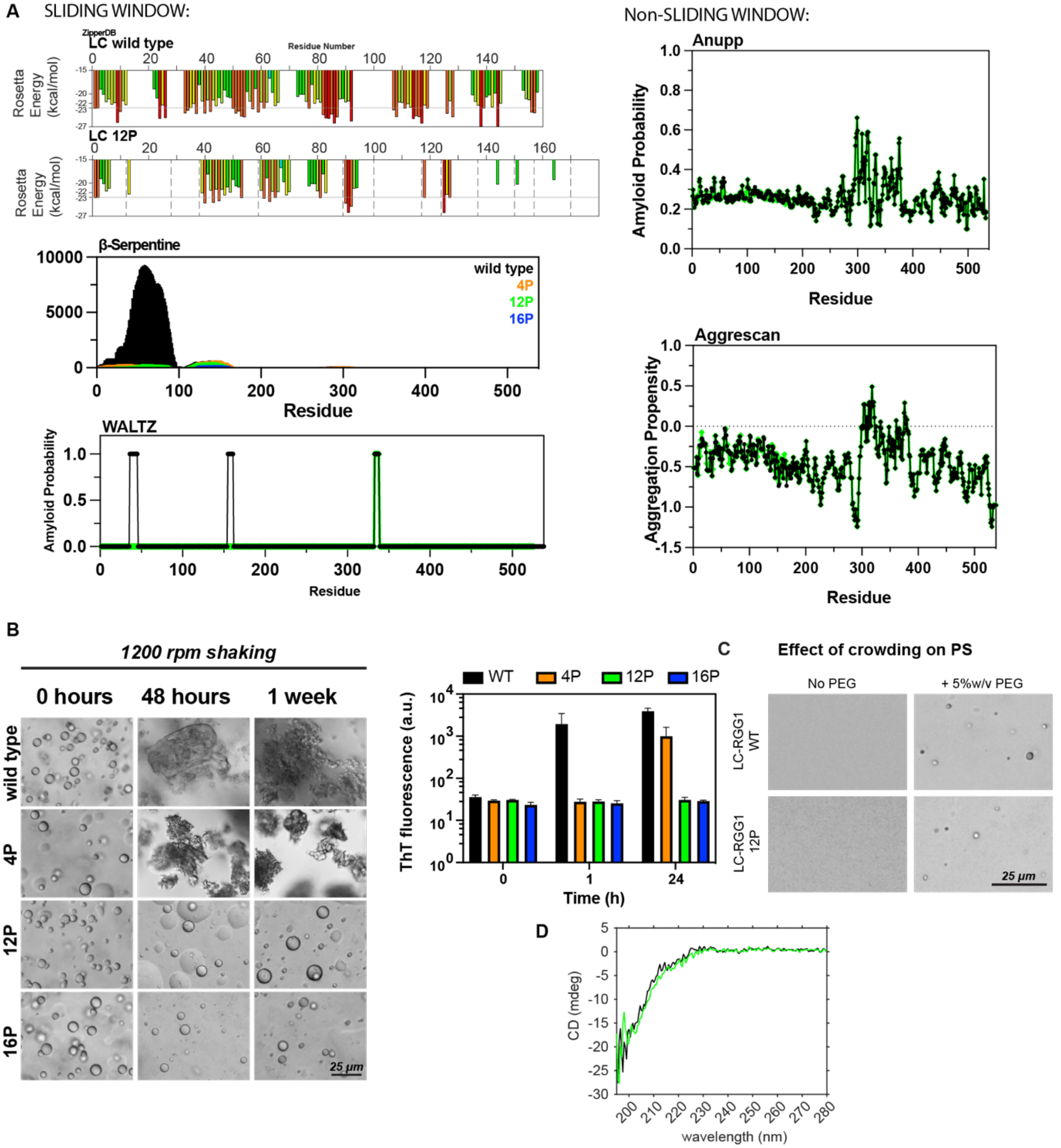


**Supplementary Figure 1. Design and analysis of proline insertion variants**

**A)** Sliding-window aggregation predictors predict disruption of β-sheet formation to different extents whereas algorithms that do not consider local sequence context show little difference with proline variants.

**B)** (Left) DIC micrographs of FUS LC variants 4P, 12P, and 16P exposed to increasing durations of 1200 rpm orbital shaking. The 12P and 16P variants show resistance to aggregation for up to 1 week. (Right) Fluorescence of Thioflavin T was measured for samples 300 μM FUS LC shaken at 1200 rpm for up to 24 hours. Bars represent the average of n = 3 technical replicates.

**C)** DIC micrographs of 5 μM FUS LC-RGG1 wild type and 12P in the presence of 5% w/v PEG.

**D)** Concentration of the condensed phase was directly measured for the FUS LC-RGG1 wild-type and 12P.

**E)** Circular dichroism spectra of the FUS LC wild-type or 12P in the dispersed phase. Samples were prepared as 5 μM FUS LC in 20 mM sodium phosphate, pH 7.0, 150 mM sodium chloride. Data are presented as mean of n = 3 technical replicates.

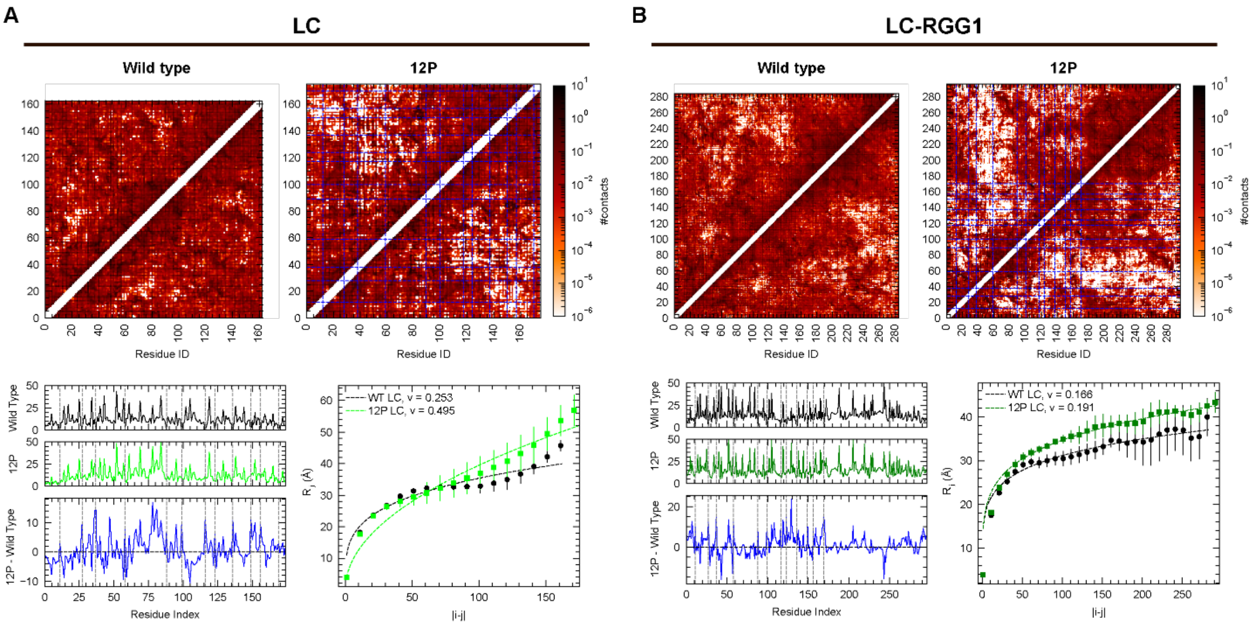


**Supplementary Figure 2: Impact of 12P mutations on the conformational ensembles of LC and LC-RGG1.**

**A)** Analysis of the isolated LC domain and **(B)** the LC-RGG1 construct. **Top**: Intramolecular simulation contact maps for wild-type and 12P variants show a reduction in long-range contacts in the 12P mutants. **Bottom Left**: Per-residue contact number for wild-type and 12P, alongside the difference profile. **Bottom Right**: The average intrachain distance $R_{ij}$ between residues i and j calculated from simulations. Error bars represent the s.e.m. from three independent trajectories. Dashed lines indicate best fits to the power law $R_{ij}=|i\text{-}j|^{v}$, where $v$ is the scaling exponent.

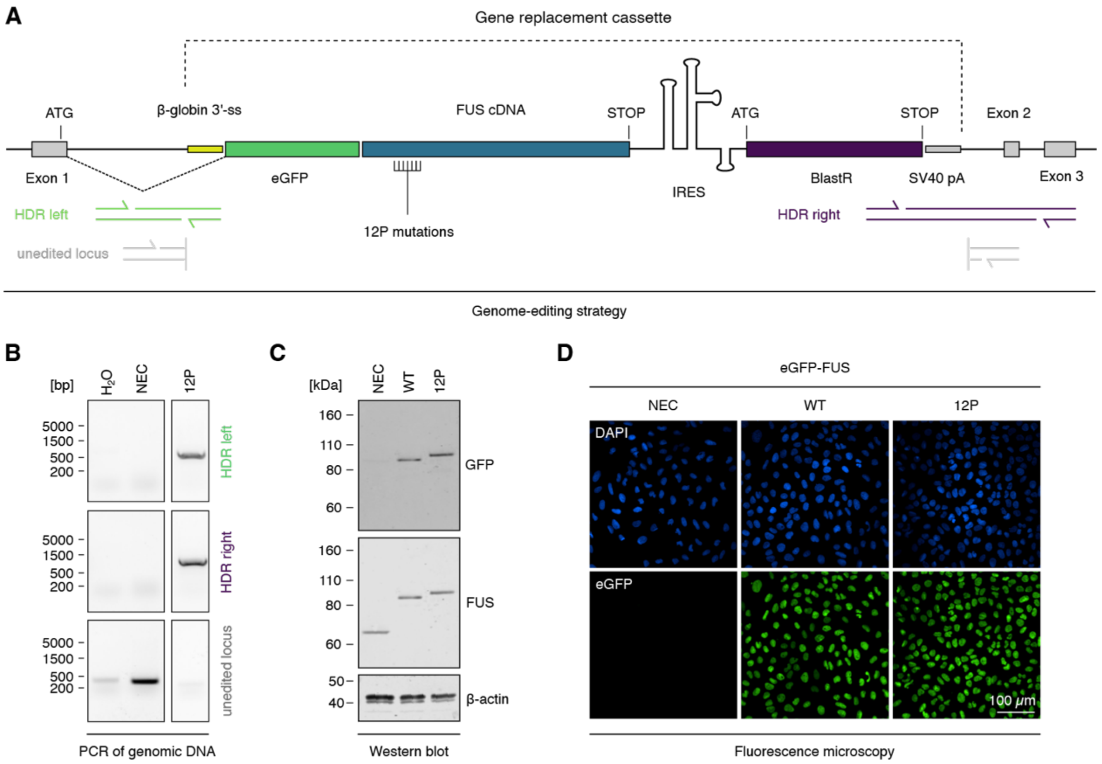


**Supplementary Figure 3**. **Design and validation of nuclear eGFP-FUS U2OS cell lines.**

**A)** Schematic depiction of the genome-editing approach used to knock-in eGFP-FUS cDNA into the first intron of the endogenous *FUS* locus and binding sites of primers used for genotyping by PCR across homology-directed repair (HDR) junctions.

**B)** Agarose gel confirming homozygous integration of the cDNA cassette into the endogenous *FUS* locus.

**C)** Western blot analysis of eGFP-FUS constructs expression using anti-GFP and anti-FUS antibodies. β-Actin was used as a loading control. 12P eGFP-FUS constructs show a mobility shift due to the 12 proline residues addition, confirming mutant identity.

**D)** Fluorescence microscopy images of eGFP-FUS (green) and DAPI (blue) in edited cell lines showing purity and homogeneous nuclear FUS expression. Scale bar, 100 µm (nuclear lines).

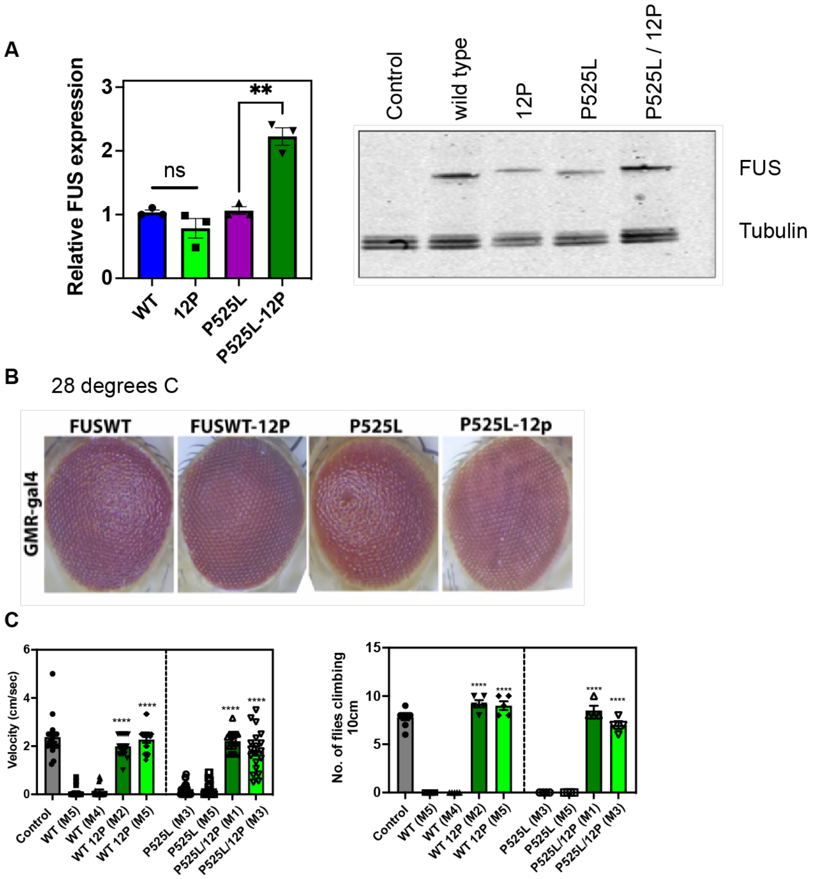


**Supplementary Figure 4.**

**A)** Quantification of Western Blot measuring total expression levels of FUS variants in *Drosophila* model.

**B)** GMR-gal4 expression of FUS variants at 28 °C using an eye-specific driver in *Drosophila*.

**C)** Further quantification of fly motility and their ability to migrate 5 and 10 cm, and velocity quantification.

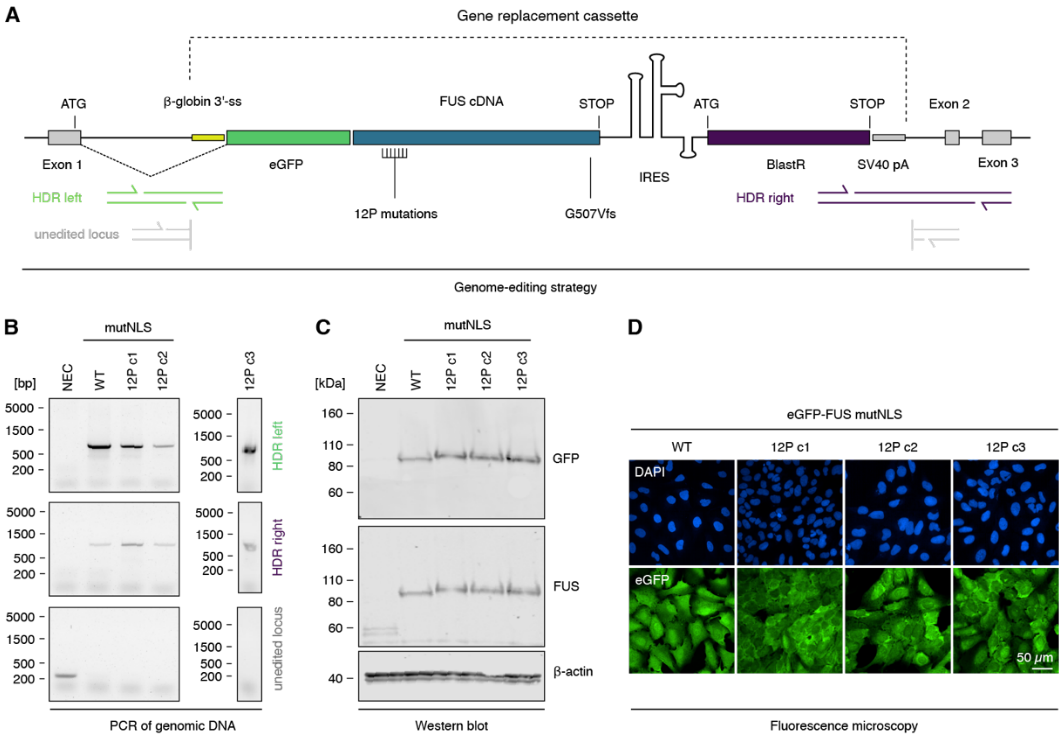


**Supplementary Figure 5. Design and validation of cytoplasmic eGFP-FUS U2OS cell lines.**

**A)** Schematic depiction of the genome-editing approach for the cytoplasmic FUS lines.

**B)** Agarose gel confirming homozygous integration of the cDNA cassette into the endogenous *FUS* locus.

**C)** Western blot analysis of cytoplasmic eGFP-FUS and three different clones of cytoplasmic 12P eGFP-FUS construct expression using anti-GFP and anti-FUS antibodies. β-Actin was used as a loading control.

**D)** Fluorescence microscopy images of cytoplasmic eGFP-FUS and 12P eGFP-FUS clones (green) and DAPI (blue) in edited cell lines showing purity and homogeneous cytoplasmic FUS expression. Scale bar, 50 μm.

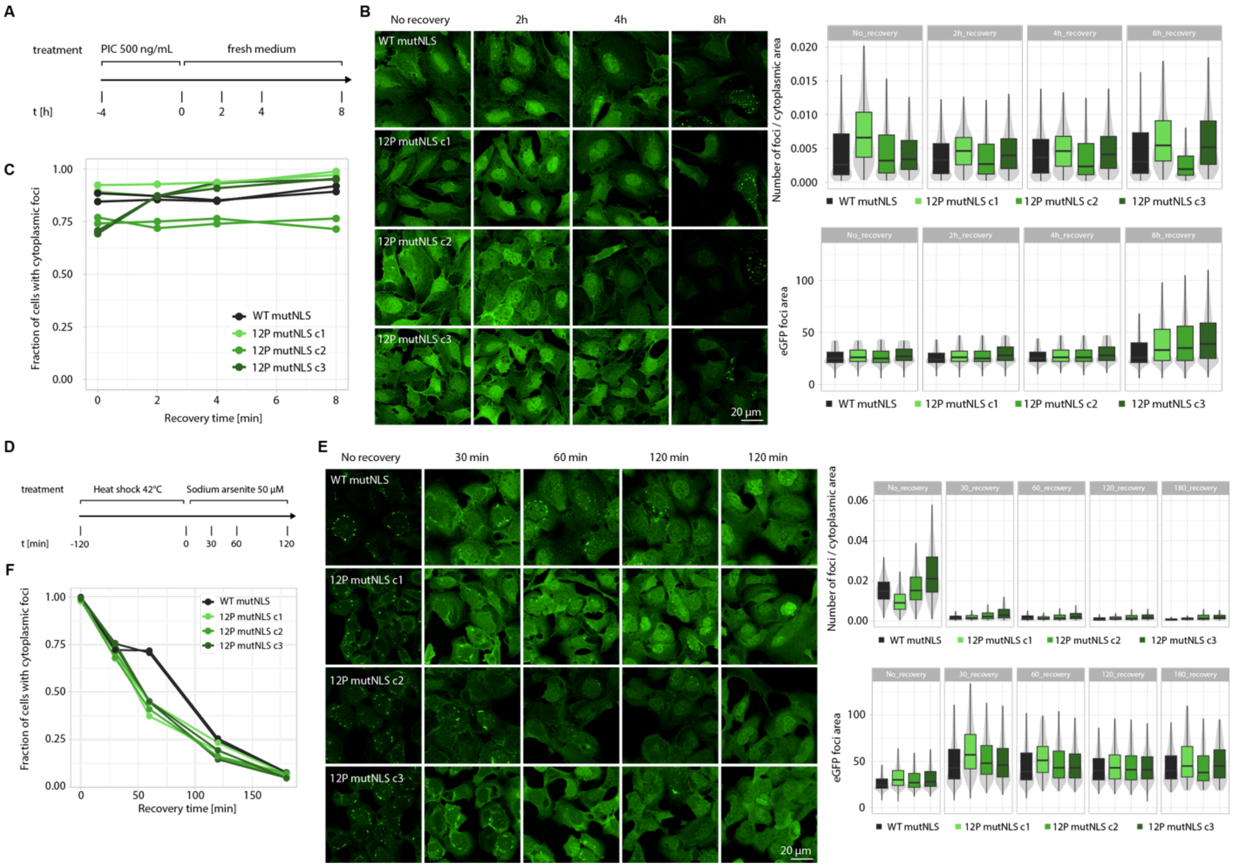


**Supplementary Figure 6. Testing the impact of β-sheet disrupting proline additions on cytoplasmic FUS accumulation in poly(I:C)-induced stress granules**

We sought to test whether stress condensates that were formed in response to viral nucleic acid showed evidence of condensate maturation. Motivated by studies from Shelkovnikova et al.[1] we exposed U2OS cells to poly(I:C) viral mimic dsRNA synthetic analog for 4h, followed by a wash-off and time-lapse tracking of granule formation and recovery; however, we were unable to reproduce the finding of persistent FUS-containing stress granules in the cytoplasm. Because it has been reported that different lengths of poly(I:C) nucleic acid can elicit different cellular responses, we used both low-molecular-weight (LMW) (#tlrl-picw) (**A,B,C**) and high-molecular-weight (HMW) (#P9582, not shown) poly(I:C) segments. We found that after 4h treatment, only ≥ 500 ng/µl LMW poly(I:C) triggered a response in our U2OS cell lines expressing cytoplasmic FUS (**Fig. S5**), but still were not successful in forming persistent stress granules. Alternatively, we found that exposing U2OS cells to brief heat shock followed by low-level sustained sodium arsenite treatment created a minor population of persistent structures for the wild type (**D, E**). While, compared with the wild type, the cells expressing the 12P variant did not form this population of persistent assemblies—suggesting a modest increase in condensate reversibility — it remains unclear how consistently this effect translates across different cell lines. Therefore, this finding should be carefully interpreted, but is included here to provide information regarding the different responses to poly(I:C) in the U2OS cell line as has been reported elsewhere[2-4].

1. Shelkovnikova, T.A. et al. Antiviral Immune Response as a Trigger of FUS Proteinopathy in Amyotrophic Lateral Sclerosis. *Cell Rep* **29**, 4496-4508 e4 (2019).
2. Shang, Z. et al. TRIM25 predominately associates with anti-viral stress granules. *Nat Commun* **15**, 4127 (2024).
3. Manjunath, L. et al. APOBEC3B drives PKR-mediated translation shutdown and protects stress granules in response to viral infection. *Nat Commun* **14**, 820 (2023).
4. Yoshioka, D., Nakamura, T., Kubota, Y. & Takekawa, M. Formation of the NLRP3 inflammasome inhibits stress granule assembly by multiple mechanisms. *J Biochem* **175**, 629-641 (2024).
